# Physical exercise restores adult neurogenesis deficits induced by simulated microgravity

**DOI:** 10.1101/2023.10.27.564332

**Authors:** Alexandra Gros, Fandilla Marie Furlan, Vanessa Rouglan, Alexandre Favereaux, Bruno Bontempi, Jean-Luc Morel

## Abstract

Cognitive impairments have been reported in astronauts during spaceflights and documented in ground-based models of simulated microgravity (SMG) in animals. However, the neuronal causes of these behavioral effects remain largely unknown. We explored whether adult neurogenesis, known to be a crucial plasticity mechanism supporting memory processes, is altered by SMG. Adult male Long-Evans rats were submitted to the hindlimb unloading model of SMG. We studied the proliferation, survival and maturation of newborn cells in the following neurogenic niches: the subventricular zone (SVZ)/olfactory bulb (OB) and the dentate gyrus (DG) of the hippocampus, at different delays following various periods of SMG. SMG exposure for 7 days, but not shorter periods of 6 or 24 hours, resulted in a decrease of newborn cell proliferation restricted to the DG. SMG also induced a decrease in short-term (7 days), but not long-term (21 days), survival of newborn cells in the SVZ/OB and DG. Physical exercise, used as a countermeasure, was able to reverse the decrease in newborn cell survival observed in the SVZ and DG. In addition, depending on the duration of SMG periods, transcriptomic analysis revealed modifications in gene expression involved in neurogenesis. These findings highlight the sensitivity of adult neurogenesis to gravitational environmental factors during a transient period, suggesting that there is a period of adaptation of physiological systems to this new environment.

## Introduction

The increased duration and distance of future space missions may compromise the health of crew-members who will have to adapt to a highly challenging environment. Numerous environmental factors constitute stressors for astronauts: microgravity and altered gravity, exposure to ionizing particle radiation, social isolation, and other spaceflight hazards (Afshinnekoo et al., 2020). These stressful stimuli could affect physiological systems, triggering biological adaptive changes to re-establish the optimal homeostatic state. During spaceflights, astronauts report mental fatigue, disruptions of neurovestibular system, sleep and circadian rhythms, and difficulty concentrating and maintaining a high level of cognitive performance (Arshad and Ferré, 2023; Gupta et al., 2023; Mhatre et al., 2022; Oluwafemi et al., 2021). MRI investigations have revealed that the brain of astronauts undergoes a variety of changes as a consequence of spaceflight missions including changes in tissue volume and microstructure, cerebrospinal fluid distribution and dynamics, and functional connectivity (Gupta et al., 2023; Jillings et al., 2023; Roy-O’Reilly et al., 2021). Understanding how long-term exposure to reduced-gravity environments may influence the neurophysiological processes and neuroplasticity, *i.e.*, the ability of the brain to organize its structure and related functions in response to stressful environmental stimuli has crucial translational significance for the health and adaptation of astronauts.

Animal model research is essential to identifying and understanding the mechanisms underlying space-related neurobiological and behavioral changes that could affect mental health and performance of astronauts. Rodent models have been developed in laboratories for years, in particular, the hindlimb unloading (HU) model, which simulates microgravity on Earth (Morey-Holton and Globus, 2002). This model reproduces in rodents the principal effects of microgravity observed in humans such as atrophy in unloaded muscles, bone mineralization loss (David et al., 2006; Doty et al., 2005), and cardiovascular deconditioning (Globus and Morey-Holton, 2016). Studies using HU models have shown that exposure to simulated microgravity (SMG) can cause modifications in neurotransmission (D’Amelio et al., 1998, 1996; Feng et al., 2016a; Kokhan et al., 2017; Shang et al., 2017a; Wang et al., 2017a, 2015; Wu et al., 2017a), and monoamine distribution (Gros et al., 2021) as well as morphological and electrophysiological changes in hippocampal neurons (Ranjan et al., 2014; Wang et al., 2020; Wu et al., 2017; Xiang et al., 2019; Zhai et al., 2019). Moreover, modifications in protein and gene expression in the hippocampus and cerebral cortex have been described (Chen et al., 2016; Frigeri et al., 2008; Malkani et al., 2020; Overbey et al., 2019; Wang et al., 2017; Wu et al., 2017; Xiang et al., 2019; Zhai et al., 2019) and suggested to underlie cognitive deficits (Feng et al., 2016a; Kokhan et al., 2017; Shang et al., 2017a; Wang et al., 2020a, 2017a; Wu et al., 2017a; Xiang et al., 2019a; Zhai et al., 2019).

Learning and memory processes are supported by neuronal plasticity mechanisms, including adult neurogenesis (Gros et al., 2015). In the mammalian brain, new neurons are continuously generated throughout life in restricted brain areas called neurogenic niches: the sub-granular zone of the dentate gyrus (DG) of the hippocampus where new dentate granule cells are generated, and the sub-ventricular zone (SVZ) where new neurons are generated to supply the olfactory bulb (OB) in interneurons (Ming and Song, 2011; Zhao et al., 2008). The process of adult neurogenesis starts with neural progenitor cells or stem cells. These progenitor cells divide many times before specializing into different cell types, mostly neurons. Adult neurogenesis involves an initial period of neural proliferation, followed by a period of survival, selection and maturation over the course of several weeks (Ming and Song, 2011; Zhao et al., 2008). This dynamic and finely-regulated process is highly dependent on the activity of neural networks and subject to modulation by various physiological and environmental stimuli (Aimone et al., 2014). The impact of spaceflight on adult neurogenesis has been little explored. It is well-known that irradiation impairs adult neurogenesis in the SVZ and DG (Madsen et al., 2003; Nagai et al., 2000; Peißner et al., 1999; Rola et al., 2004; Tada et al., 2000), making it a standard uninvasive method to delete new neurons in specific neurogenic regions of adult rodents (Wojtowicz, 2006). Social isolation and/or exposure to an impoverished environment are also known to impair adult neurogenesis (Grégoire et al., 2014; Holmes, 2016; Ibi et al., 2008; Lu et al., 2003). However, the impact of microgravity on adult neurogenesis remains understudied. *In* vitro, neural stem cells derived from the SVZ of HU mice present a reduced proliferation capability, an altered cell cycle and an incomplete differentiation/maturation (Adami et al., 2018). *In* vivo, HU for 2 weeks reduces the number of proliferating cells in the SVZ (Adami et al., 2018; Yasuhara et al., 2007) and DG (Yasuhara et al., 2007). However, these studies were limited to proliferation and did not explore newborn cell survival and maturation.

Here, we explored the impact of SMG on the different stages of the adult neurogenesis process in male Long Evans rats experiencing HU. Moreover, we evaluated the role of physical exercise as a countermeasure to protect or limit the effects of SMG on adult neurogenesis. Physical exercise is indeed commonly used to combat the deleterious physiological effects of microgravity exposure in astronauts during spaceflights (English et al., 2020; Loehr et al., 2015; Scott et al., 2023). In addition, several studies demonstrated the significant positive impact of physical exercise on adult neurogenesis (Fabel and Kempermann, 2008; Vivar et al., 2013; Voss et al., 2013). In this study, we found that SMG impaired newborn cell proliferation in the DG only after 7 days of exposure. SMG also decreased the short-term (7 days), but not long-term (21 days) survival of newborn cells in both the SVZ/OB and the DG. Physical exercise was able to reverse the impact of SMG on newborn survival observed in these two areas. At the molecular level, RTqPCR and RNA-seq analyses revealed a transient change in the expression of several genes involved in neurogenesis. These findings suggest that adult neurogenesis is transiently impaired by SMG and identify physical exercise as an effective countermeasure to limit the impact of SMG on new neurons generated throughout adulthood.

## Results

### Validation of the HU model

In this study, the effects of microgravity were mimicked by submitting male Long Evans rats to the HU model (Figure 1A). To assess its efficiency in our experimental conditions, we weighed the soleus muscle which controls the hindlimbs and known to be impacted by microgravity in astronauts and in the HU model (Ohira, 2000). The extensor carpi radialis longus (ECRL) muscle in the forward limbs, whose weight is not impacted by microgravity in astronauts and in the HU model, served as control. As expected, 7 days of SMG exposure significantly decreased the weight of the soleus muscle (Figure 1B; Unpaired t-test, t_6_ = 6.57, p < 0.001), but not the weight of the ECRL muscle (Figure 1C; Unpaired t-test, t_6_ = 0.07, p = 0.95). The same pattern of effects was observed when the weight of these two muscles was expressed as a function of the total weight of the rats (Figure Sup. 1E). Thus, these results validate the effects of HU in our experimental context.

**Figure 1.**
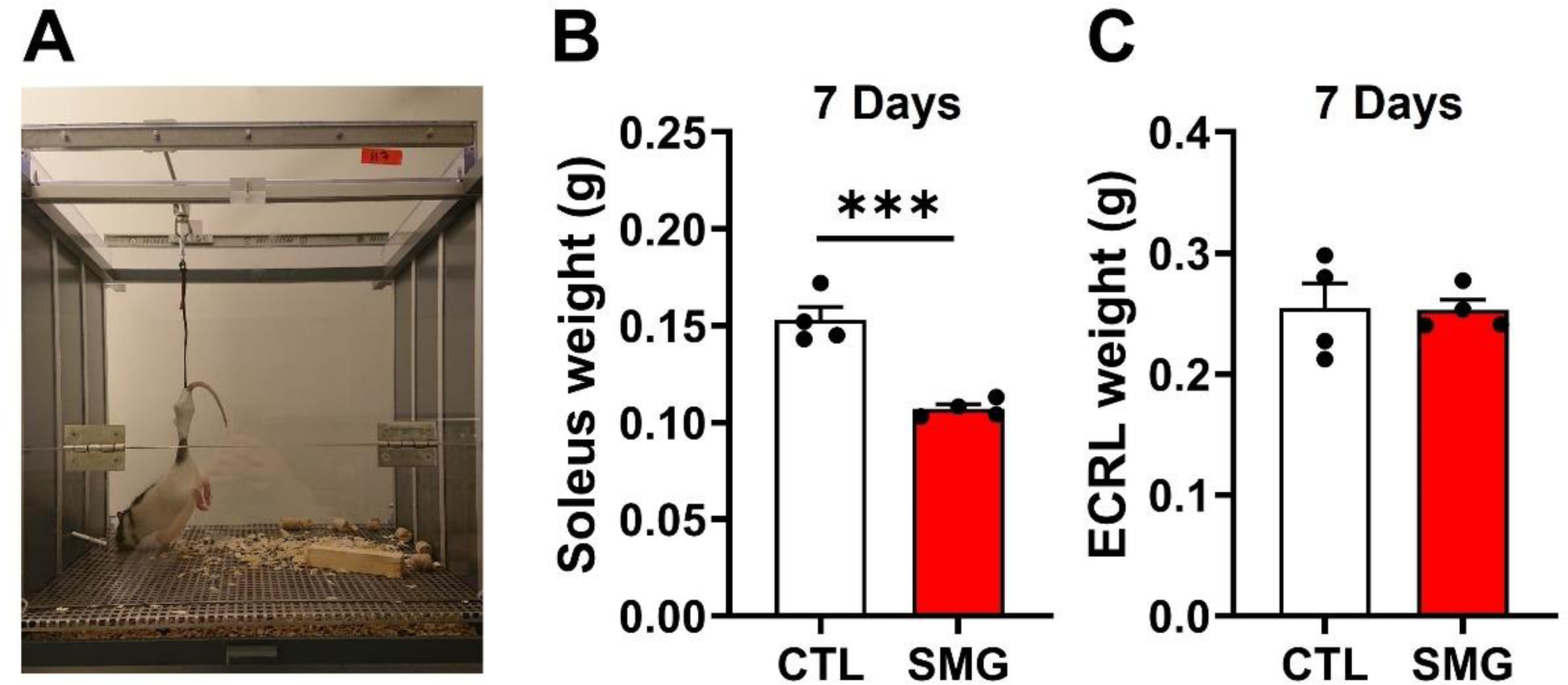
Validation of HU model. **A.** Picture of the HU model. **B and C.** Effects of SMG on the weight of the soleus (B) and extensor carpi radialis longus (ECRL) (B) muscles in CTL (white) and SMG (red) rats. All data are presented as mean ± SEM. *** p < 0.005.

### Adult neurogenesis

#### Simulated microgravity alters the weight gain in rats injected with adult neurogenesis marker

The physiological parameters of the rats were carefully monitored during the whole period of SMG exposure. To study adult neurogenesis, rats were injected with the cell birth marker EdU (Day 0) and immediately exposed to hindlimb suspension (SMG) or kept in control condition (CTL) for different time periods ranging from 6h to 21 days (Figure 2A). EdU injections induced a stress in both CTL and SMG rats as revealed by a high concentration of corticosterone in plasma 6h after EdU administration (Figure 2B). No difference was observed between CTL and SMG rats (two-way ANOVA, SMG effect F(1, 52) = 0.54, p = 0.47). Moreover, we observed a significant time effect (two-way ANOVA, time effect F(3, 52) = 6,89, p < 0.001). However, corticosterone concentration significantly decreased over time in CTL (white bars; one-way ANOVA F(3, 26) = 6.59, p = 0.002) but not in SMG rats (red bars; one-way ANOVA F(3, 26) = 2.01, p = 0.14), suggesting that the injection-related stress was potentiated in the SMG condition.

**Figure 2.**
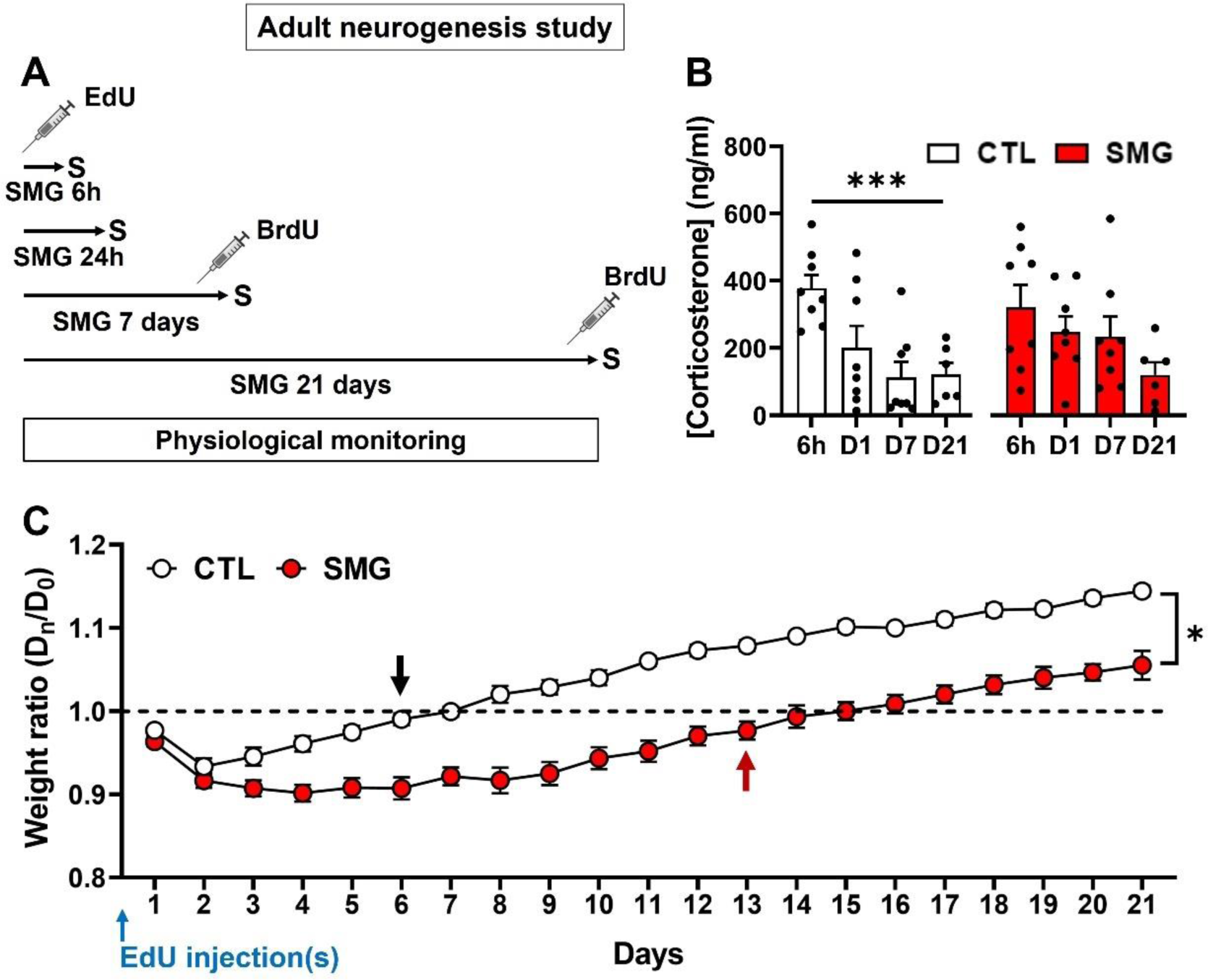
SMG caused a delay in the weight gain of rats injected with the adult neurogenesis marker. **A.** Experimental design. EdU was injected just before hindlimb suspension. Rats were exposed to SMG during 6 hours, 24 hours, 7 days or 21 days. Rats exposed to 7 or 21 days were additionally injected with BrdU 24h before sacrifice (S). The physiological parameters of the animals were monitored daily for the entire duration of the protocol. **B.** Corticosterone concentration in CTL (white) and SMG (red) rats. **C.** Body weight of the CTL (white) and SMG (red) rats presented as a ratio relative to their weight at the day of Edu injections. The black and red arrows indicate a return of CTL and SMG rats to their starting weight, respectively. All data are presented as mean ± SEM. * p < 0.05, *** p < 0.005.

The body weight of CTL rats was decreased compared to their starting weight after EdU administration during 5 days (Figure 2C, white circles; one-sample t-test, D1 to D5 p < 0.005). These rats required 6 days to return to their starting body weight (black arrow; one-sample t-test, D6 to D8 p > 0.1) and 9 days to significantly gain weight relative to D0 (one-sample t-test, D9 to D21 p < 0.01). The body weight of rats exposed to SMG was also decreased by the EdU administration (Figure 2C, red circles; one-sample t-test, D1 to D7 p < 0.0001, D8 to D11 p < 0.01, D12 p = 0.045). However, it took them 13 days to return to their starting body weight (red arrow; one-sample t-test, D13 p = 0.08, D14 to D17 p > 0.1) and 18 days to significantly increase their body weight (one-sample t-test, D18 to D21 p < 0.05). A significant difference was observed between CTL and SMG rats from D4 to the end of the experiment (two-way repeated measures ANOVA, mixed-effects model, SMG effect F(1, 42) = 60.91, p < 0.001; Bonferroni multiple comparisons test, D1 to D3 p > 0.2, D4 to D21 p < 0.05). A simple linear regression over 21 days showed that weight gain differed between CTL and SMG rats (F(1, 396) = 41.98, p < 0.0001). A more detailed analysis revealed that this difference was due to the first 4 days of exposure. Indeed, from D5, the simple linear regression showed no difference between CTL and SMG rats (F(1, 236) = 0.06, p = 0.82), indicating that after an initial phase of body weight stagnation, weight gain in SMG and CTL rats was similar.

The EdU injection effect on the body weight was in part due to a decrease in food consumption observed during the first 4 days in CTL and SMG rats compared with the food consumption at D0 recorded before EdU injection(s) (Figure Sup. 1A; two-way repeated measures ANOVA, mixed-effect model, time effect F(3.92, 67.09) = 34.09, p < 0.0001; Dunnett multiple comparisons test *vs* D0, CTL: D1 to D4 p < 0.01, SMG: D1 to D4 p < 0.001). Then, food consumption was similar to the initial consumption in CTL rats (D5 to D21 p > 0.1) but more variable in SMG rats (D5 to D7 p > 0.05, D8 p = 0.044, D9 to D19 p > 0.1, D20 p = 0.027, D21 p = 0.99). However, no difference was observed between CTL and SMG rats at any time (two-way repeated measures ANOVA, mixed-effects model, SMG effect F(1, 43) = 5.18, p = 0.028; Bonferroni multiple comparisons test, D1 to D21 p > 0.25). This result indicates that despite SMG rats consuming the same amount of food as CTL rats, their weight gain decreased. Water consumption was also slightly altered the first day following the EdU injection(s) (Figure Sup. 1B; two-way repeated measures ANOVA, mixed-effect model, time effect F(5.57, 92.76) = 7.26, p < 0.0001; Dunnett multiple comparisons test *vs* D0, CTL: D1 p = 0.0009, D2 to D21 p > 0.15; SMG: D1 p < 0.0001; D2 to D21 p > 0.15). As for food consumption, no difference was observed between CTL and SMG rats (two-way repeated measures ANOVA, mixed-effects model, SMG effect F(1, 43) = 0.022, p = 0.88). No effect of SMG was observed in rat daily temperature (Figure Sup. 1C; two-way repeated measures ANOVA, mixed-effect model, time effect F(1, 43) = 2.32, p = 0.14) or in glycemia measured at D0, D1, D7, D14 and D21 (Figure Sup. 1D; two-way repeated measures ANOVA, mixed-effect model, time effect F(1, 43) = 0.08, p = 0.79). Both temperature and glycemia measures were within physiological values (green area).

Overall, SMG potentiated the stressful effect of the EdU injection(s) notably with an effect on the rats’ body weight.

#### Simulated microgravity transiently alters the proliferation of newborn hippocampal cells

We evaluated the impact of SMG on newborn cell proliferation in the DG and SVZ of rats injected with EdU and then immediately exposed to SMG during 24h (Figure 3A and B). SMG did not affect EdU cell density in the DG (Figure 3C; Unpaired t-test, t_9_ = 0.52, p = 0.61) or in the SVZ (Figure 3D; Mann-Whitney test, p = 0.54). Similar results were observed after 6 hours of SMG exposure (Figure Sup. 2A-C; DG: unpaired t-test, t_9_ = 0.37, p = 0.72; SVZ: unpaired t-test, t_9_ = 1.59, p = 0.15). As expected, we observed the well-known time-dependent increase in newborn cell proliferation in the DG between 6 and 24h (Figure Sup. 2B; two-way ANOVA, time effect F(1, 18) = 8.60, p = 0.009). Moreover, the expression of GFAP (Figure Sup. 2D) and Sox2 (Figure Sup. 2E), known to be expressed in newborn cell during the first step of proliferation, were not modified by the SMG (Figure Sup. 2F-G; three-way ANOVA, SMG effect, DG: F(1, 36) = 0.06, p = 0.8; SVZ: F(1, 36) = 0.6, p = 0.45). These results indicate that short SMG exposure did not disturb proliferation of the newborn cells in the neurogenic niche.

**Figure 3.**
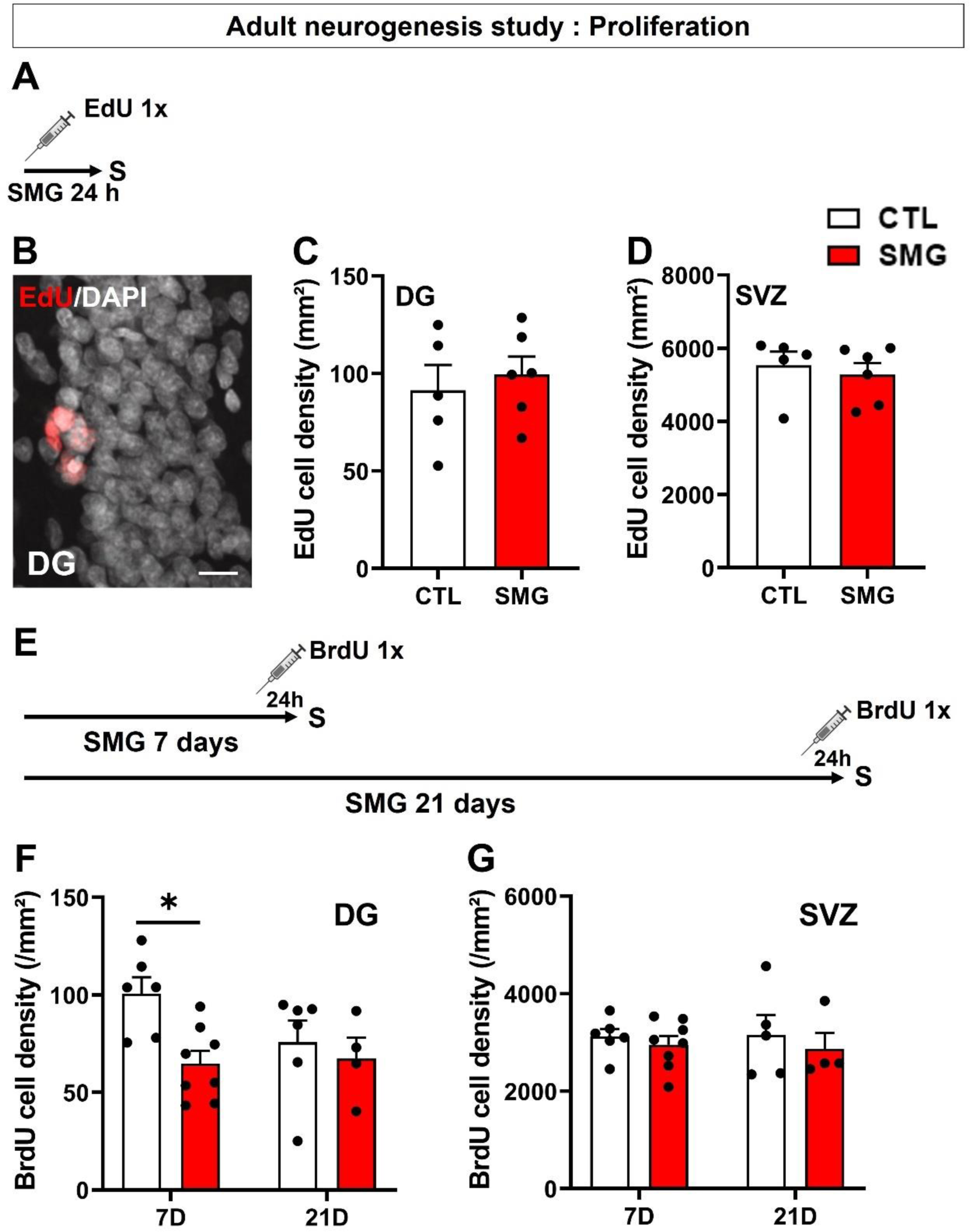
SMG impaired newborn cell proliferation after 7 days of exposure only in the DG. **A.** Experimental design. Rats were injected with EdU and exposed to SMG during 24 hours. **B.** Example of EdU+ cells (in red) in the DG of the hippocampus. Scale bar: 10 µm. **C.** Density of EdU cells per mm² in the DG of CTL (white) and SMG (red) rats. **D.** Density of EdU cells per mm² in the SVZ of CTL (white) and SMG (red) rats. **E.** Experimental design. Rats were exposed to SMG during 7 or 21 days and injected with BrdU 24h before the end of the protocol. **F.** Density of BrdU cells per mm² in the DG of CTL (white) and SMG (red) rats. **G.** Density of BrdU cells per mm² in the SVZ of CTL (white) and SMG (red) rats. All data are presented as mean ± SEM. * p < 0.05.

Because the effects of microgravity can take time to develop in both humans and animals, we next explored if longer exposure to SMG could impair the proliferation phase of adult neurogenesis. To this end, we exposed rats to SMG or control condition during 7 or 21 days. BrdU was injected 24h before the end of the hindlimb suspension to specifically tag the proliferation phase (Figure 3E). A significant decrease in BrdU cell density was observed after 7 days of SMG exposure in the DG (Figure 3F; two-way ANOVA, SMG effect F(1, 20) = 5.76, p = 0.026; Bonferroni multiple comparisons test p = 0.013), but not after 21 days (Bonferroni multiple comparisons test p > 0.99). No effect was observed in the SVZ (Figure 3G; two-way ANOVA, SMG effect F(1, 19) = 0.75, p = 0.4). These results indicate that longer exposure to SMG transiently disturbs hippocampal newborn cell proliferation.

#### Simulated microgravity transiently alters newborn cell survival

We next explored the effect of SMG on newborn cell survival in the DG, SVZ and OB. Rats were injected with EdU and then immediately exposed to SMG during 7 or 21 days (Figure 4A). As expected, we observed in CTL rats the well-known time-dependent decrease in newborn cell survival between 7 and 21 days in the 3 structures (DG: Figure 4D, two-way ANOVA, time effect F(1, 24) = 16.9, p = 0.0004; SVZ: Figure 4F, two-way ANOVA, time effect F(1, 23) = 51.96, p < 0.001; OB: Figure 4H, two-way ANOVA, time effect F(1, 23) = 142.1, p < 0.001). In the DG, EdU cell density was significantly decreased after the 7-day SMG exposure period (Figure 4D; two-way ANOVA, SMG effect F(1, 24) = 2.17, p = 0.15, interaction F(1, 24) = 5.18, p = 0.032; Bonferroni multiple comparisons test p = 0.017) but not after 21 days (p > 0.99). A similar result was observed in the SVZ (Figure 4F; two-way ANOVA, SMG effect F(1, 23) = 6.4, p = 0.019; Bonferroni multiple comparisons tests: 7D p = 0.003; 21D p > 0.99). In addition, social isolation was not a confounding factor (Figure Sup. 3). In the OB, no difference was observed between CTL and SMG (Figure 4H; two-way ANOVA, SMG effect F(1, 23) = 1.17, p = 0.29).

**Figure 4.**
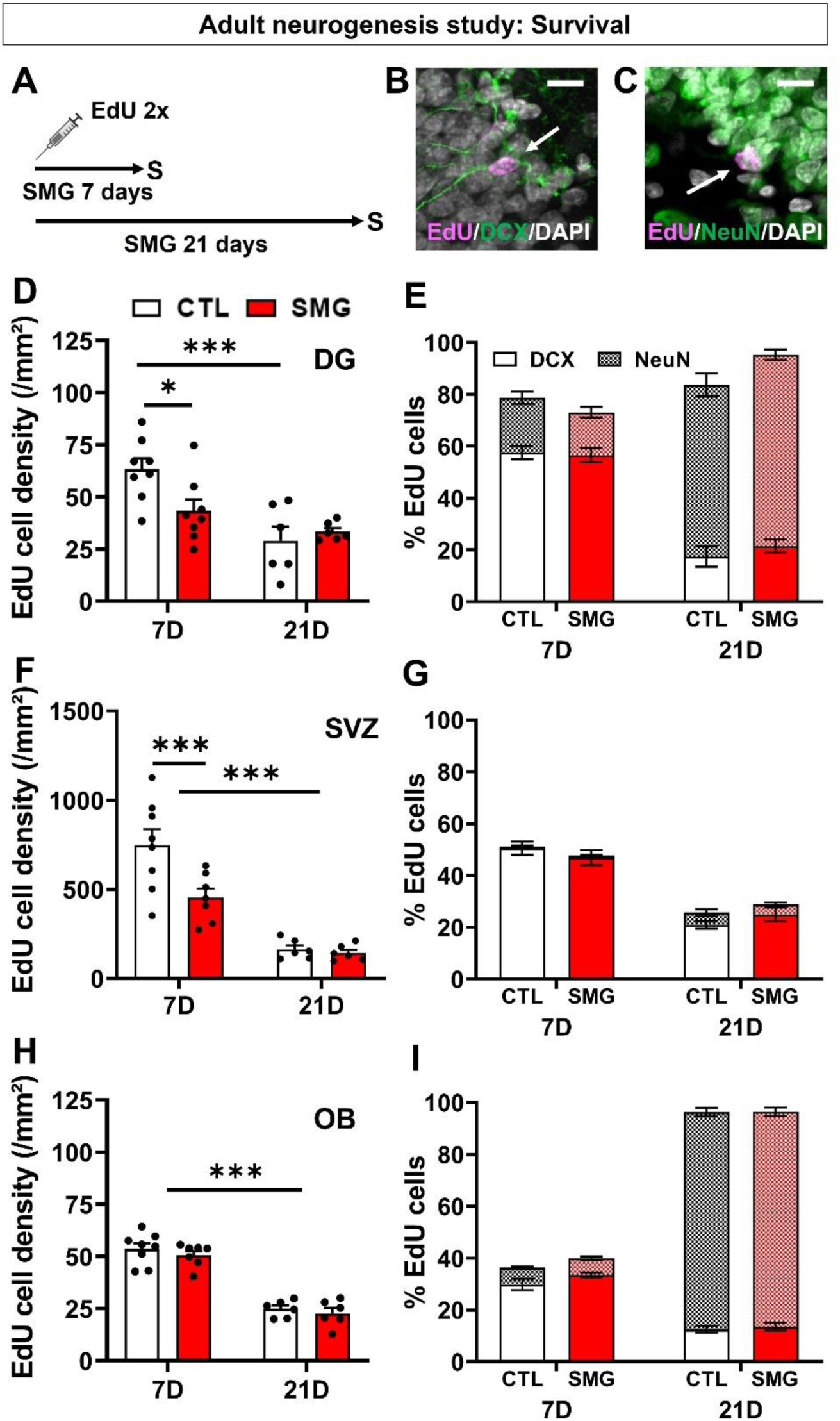
SMG resulted in impaired newborn cell survival after 7 days of exposure in the DG and SVZ. **A.** Experimental design. Rats were injected with EdU and exposed to SMG during 7 or 21 days. **B.** Example of EdU+ cell expressing DCX in the DG of the hippocampus. Scale bar: 10 µm. **C.** Example of EdU+ cell expressing NeuN in the DG of the hippocampus. Scale bar: 10 µm. **D.** Density of EdU cells per mm² in the DG of CTL (white) and SMG (red) rats. **E.** Percentage of EdU cells expressing DCX or NeuN in the DG of CTL (white) and SMG (red) rats. **F.** Density of EdU cells per mm² in the SVZ of CTL (white) and SMG (red) rats. **G.** Percentage of EdU cells expressing DCX or NeuN in the SVZ of CTL (white) and SMG (red) rats. **H.** Density of EdU cells per mm² in the OB of CTL (white) and SMG (red) rats. **I.** Percentage of EdU cells expressing DCX or NeuN in the OB of CTL (white) and SMG (red) rats. All data are presented as mean ± SEM. * p < 0.05, ** p < 0.01, *** p < 0.005.

In CTL rats, we observed a decrease in DCX expression over time associated with an increase in NeuN expression in the DG (Figure 4E, three-way ANOVA with repeated measures, time effect F(1, 23) = 20.1, p < 0.001; Bonferroni test 7D *vs* 21D: DCX p < 0.001, NeuN p < 0.001) and in OB (Figure 4I, three-way ANOVA with repeated measures, time effect F(1, 24) = 1354, p < 0.001; Bonferroni test 7D *vs* 21D: DCX p < 0.001, NeuN p < 0.001). In SVZ, we observed a decrease of DCX expression over time (Figure 4G, three-way ANOVA with repeated measures, time effect F(1, 24) = 74.04, p < 0.001; Bonferroni test 7D *vs* 21D p < 0.001). These results indicate that the neurons matured properly in the different structures analyzed. SMG did not alter the expression of DCX and/or NeuN in newborn cells in the DG (Figure 4E; three-way ANOVA with repeated measures, SMG effect F(1, 23) = 0.85, p = 0.37), the SVZ (Figure 4G; three-way ANOVA with repeated measures, SMG effect F(1, 24) = 0.005, p = 0.95) and in the OB (Figure 4I; three-way ANOVA with repeated measures, SMG effect F(1, 24) = 1.34, p = 0.26). These results indicate that exposure to SMG transiently disturb newborn cell survival in DG and SVZ, without affecting the maturation of these cells.

#### Physical exercise facilitates physical recovery after injections of the adult neurogenesis marker

During spaceflights, astronauts are submitted to a strict physical exercise routine to limit muscle loss in the forward limbs and the effects of microgravity on the cardiovascular system. We explored the impact of physical exercise on adult neurogenesis during SMG exposure. For this, rats were exposed to physical exercise in a treadmill during 2 weeks prior to the SMG exposure and 1 week during the SMG exposure (Figure 5A). Rats were injected with EdU just before the SMG exposure (Figure 5A). No difference was observed in the distance covered by the rats during the 3 weeks of exposure to physical exercise between the CTL and the SMG conditions (CTL: 461 ± 9 m/rat/day *vs* SMG: 465 ± 5 m/rat/day; unpaired t-test, t_18_ = 0.44, p = 0.67) or during the last week, i.e., during SMG exposure (CTL: 477 ± 23 m/rat/day *vs* SMG: 471 ± 15 m/rat/day, unpaired t-test, t_18_ = 0.23, p = 0.82). This indicates that SMG did not impair the motor skills of the rats to perform physical exercise on the treadmill.

**Figure 5.**
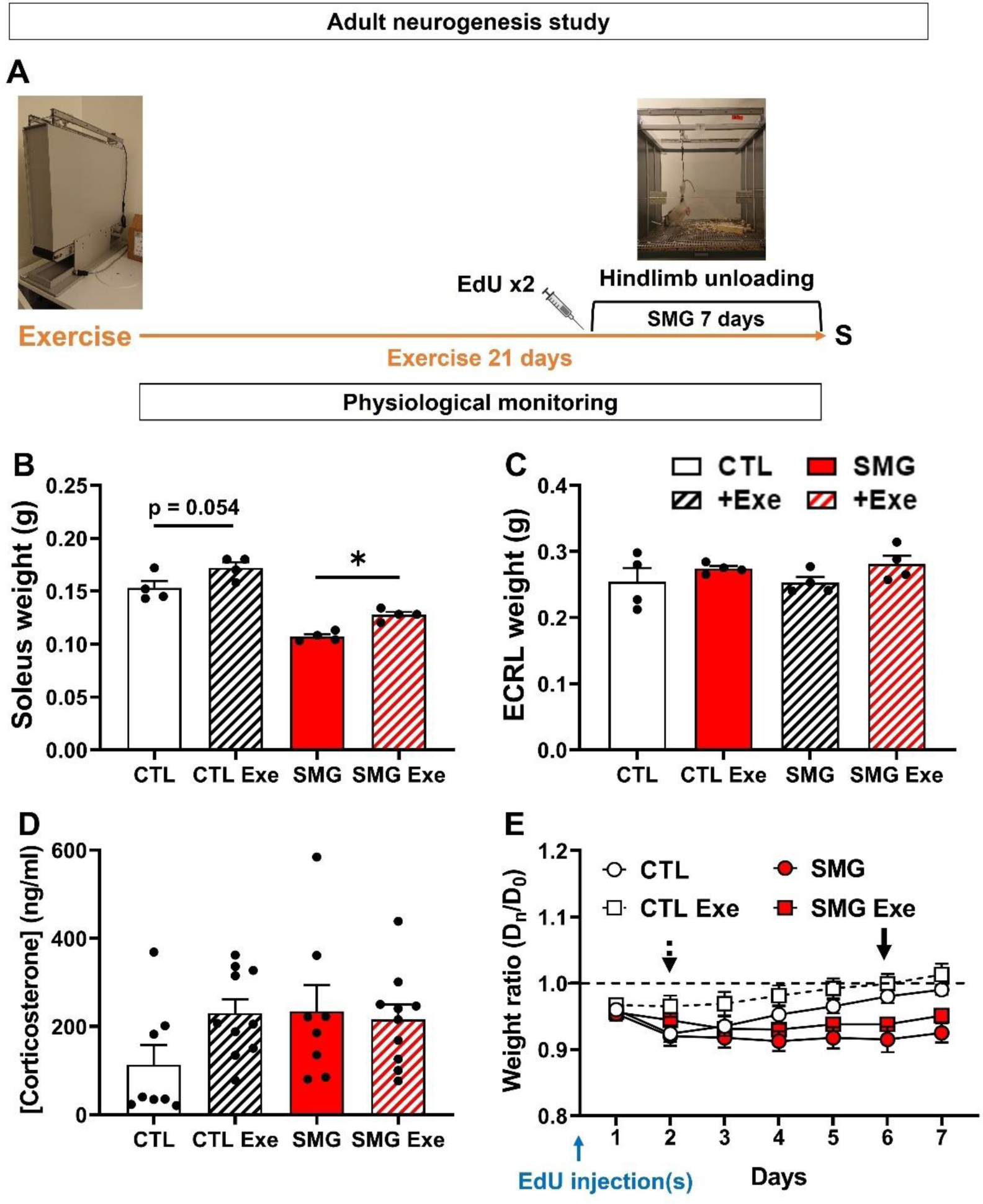
Exercise improves weight recovery in rats after injections with the adult neurogenesis marker. **A.** Experimental design. Rats were exposed to physical exercise during 3 weeks. After 2 weeks, rats were injected with EdU and exposed to SMG for 7 days. The physiological parameters of the animals were monitored daily. **B.** Weight of the soleus muscle in CTL (white) and SMG (red) rats exposed or not to physical exercise. **C.** Weight of the extensor carpi radialis longus (ECRL) muscle in CTL (white) and SMG (red) rats exposed or not to physical exercise. **D.** Corticosterone concentration in CTL (white) and SMG (red) rats exposed or not to physical exercise. **E.** Weight of the CTL (white) and SMG (red) rats exposed or not to physical exercise presented as a ratio compared with the weight at D0. Black arrow marks the point where the CTL rats return to their starting weight. The black dotted arrow marks the point where the CTL rats exposed to physical exercise return to their starting weight. All data are presented as mean ± SEM. * p < 0.05.

First, physical exercise increased the weight of the soleus muscle in CTL and SMG rats (Figure 5B; two-way ANOVA, exercise effect F(1, 12) = 18.45, p = 0.001; Tukey multiple comparisons test, CTL *vs* CTL Exe p = 0.054, SMG *vs* SMG Exe p = 0.036). No weight change was observed for the ECRL muscle (Figure 5C; two-way ANOVA, exercise effect F(1, 12) = 3.39, p = 0.09). The same result was observed when the weight of the muscles was expressed as the total body weight of the rats (Figure Sup. 4E-F). These results indicate that physical exercise is efficient in limiting muscle loss in SMG rats. Moreover, physical exercise did not change the stress level in CTL and SMG rats (Figure 5D; two-way ANOVA, exercise effect F(1, 28) = 0.89, p = 0.35).

We previously reported that the body weight of the CTL rats was decreased after EdU injections and it took them 6 days to return to their starting weight (Figure 2D and Figure 5E, black arrow). Physical exercise reduced the recovery time after EdU injections by 4 days (Figure 5E, black dotted arrow; CTL: one-sample t-test, D1 to D5 p < 0.05, D6 p = 0.08, D7 p = 0.24; CTL Exe: one-sample t-test, D1 p = 0.005, D2 p = 0.058, D3 to D7 p > 0.1). This result indicates that physical exercise facilitates weight recovery in CTL rats after EdU injections. In SMG rats, we did not observe a beneficial effect of the physical exercise on weight recovery time during the 7 days of observation. No difference was observed between rats exposed to exercise or not in both gravity condition (three-way ANOVA, exercise effect F(1, 32) = 3.58, p = 0.07, interaction SMG x Exercise F(1, 32) = 0.11, p = 0.75). Simple linear regressions showed no difference induced by physical exercise in CTL (F(1, 122) = 0.01, p = 0.91) or SMG rats (F(1, 115) = 0.24, p = 0.63). As previously observed, a significant difference between CTL and SMG rats without exercise was observed (F(1, 108) = 6.28, p = 0.014), but also with exercise (F(1, 129) = 4.9, p = 0.028). These results indicate that the gain weight was different in SMG rats compared with CTL rats during the first week of exposure to SMG.

Food consumption was reduced during 3 days after EdU injections in CTL rats (Figure Sup. 4A; two-way ANOVA, time effect F(3.533, 49.46) = 30.68, p < 0.0001; Dunnett multiple comparisons test *vs* D0: D1 to D3 p < 0.01, D4 to D7 p > 0.2) and 2 days in SMG rats (Dunnett multiple comparisons test *vs* D0: D1 to D2 p < 0.05, D3 to D7 p > 0.1). Physical exercise decreased the recovery time in CTL rats by 2 days (Dunnett multiple comparisons test *vs* D0, D1 p< 0.001, D2 to D7 p > 0.1), but not in SMG rats (Dunnett multiple comparisons test *vs* D0: D1 to D3 p < 0.01, D4 to D7 p > 0.2). No difference was observed between rats exposed to exercise or not in both gravity condition (three-way ANOVA, exercise effect F(1, 32) = 4.06, p = 0.052; Tukey multiple comparisons test p > 0.6).

Water consumption decreased the day after the EdU injections in CTL (Figure Sup. 4B; two-way ANOVA, time effect F(5.189, 72.64) = 11.72, p < 0.001; Dunnett multiple comparisons tests *vs* D0: D1 p = 0.005, D2 to D7 p > 0.5) and SMG rats (D1 p = 0.005, D2 to D7 p > 0.35). With physical exercise, CTL rats did not show decrease in water consumption (Dunnett multiple comparisons tests *vs* D0, D1 to D7 p > 0.05). In contrast, in SMG rats, physical exercise did not change the water consumption’ decrease (Dunnett multiple comparisons tests *vs* D0, D1 p = 0.052, D2 to D7 p > 0.1). Moreover, no difference was observed between rats exposed to exercise or not in both gravity conditions (three-way ANOVA, exercise effect F(1, 32) = 1.04, p = 0.31).

Physical exercise increased the body temperature in both CTL and SMG rats (Figure Sup. 4C; three-way ANOVA, exercise effect F(1, 32) = 56.43, p < 0.001). Physical exercise also increased the glycemia (Figure Sup. 4D right; D0, two-way ANOVA, exercise effect F(1, 32) = 27.06, p < 0.001). However, this effect was only visible at the end of the exercise period while no effect was observed at D7 (sacrifice day without exercise, Figure Sup. 4D left, two-way ANOVA, exercise effect F(1, 32) = 4.04, p = 0.053). In all cases, temperature and glycemia measures stayed within physiological values (green area).

#### Physical exercise restores the survival of newborn cells in DG and SVZ

As expected, physical exercise increased the EdU density in CTL rats in the DG (Figure 6B, black; two-way ANOVA, exercise effect F(1, 28) = 16.73, p < 0.001; Bonferroni multiple comparisons test p = 0.005) and in the OB (Figure 6F, black; two-way ANOVA, exercise effect F(1, 28) = 9.05, p = 0.006; Bonferroni multiple comparisons test p = 0.002). No effect of physical exercise was observed in the SVZ (Figure 6D, black; two-way ANOVA, exercise effect F(1, 27) = 4.36, p = 0.046; Bonferroni multiple comparisons test p > 0.99). These results indicate that our physical exercise paradigm was able to enhance newborn cells’ survival.

**Figure 6.**
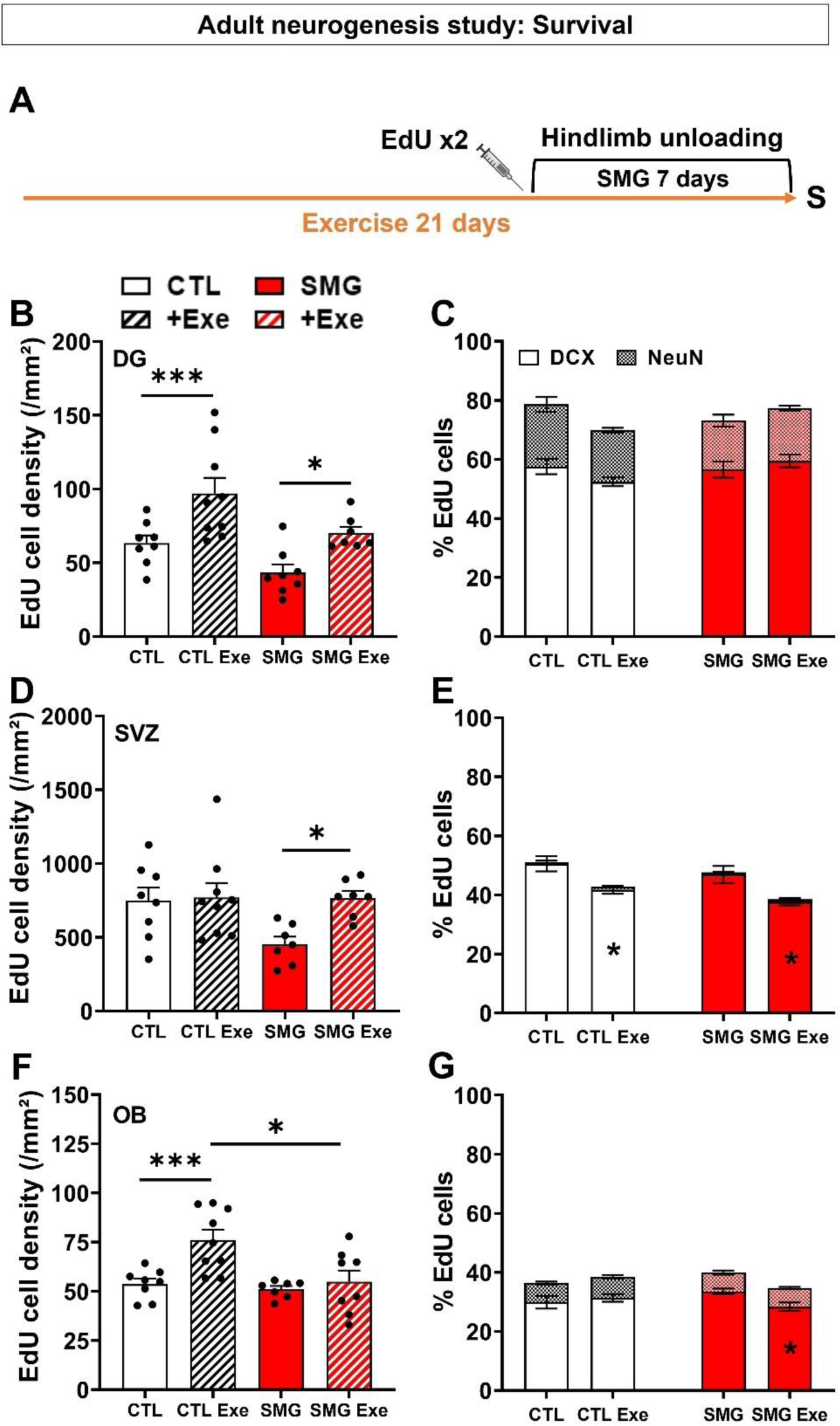
Physical exercise restored the survival of newborn cells in the DG and SVZ. **A.** Experimental design. Rats were exposed to physical exercise during 3 weeks. After 2 weeks, rats were injected with EdU and exposed to SMG for 7 days. The physiological parameters of the animals were monitored daily. **B.** Density of EdU cells per mm² in the DG of CTL (white) and SMG (red) rats exposed or not to physical exercise. **C.** Percentage of EdU cells expressing DCX or NeuN in the DG of CTL (white) and SMG (red) rats. **D.** Density of EdU cells per mm² in the SVZ of CTL (white) and SMG (red) rats exposed or not to physical exercise. **E.** Percentage of EdU cells expressing DCX or NeuN in the SVZ of CTL (white) and SMG (red) rats. **F.** Density of EdU cells per mm² in the OB of CTL (white) and SMG (red) rats exposed or not to physical exercise. **G.** Percentage of EdU cells expressing DCX or NeuN in the OB of CTL (white) and SMG (red) rats. All data are presented as mean ± SEM. * p < 0.05, *** p < 0.005.

Physical exercise increased the density of EdU cells in SMG rats in the DG (Figure 6B, Bonferroni multiple comparisons test, p = 0.038) and SVZ (Figure 6D, Bonferroni multiple comparisons test, p = 0.028). No effect of physical exercise was observed in the OB of SMG rats (Figure 6F, Bonferroni multiple comparisons test, p > 0.99). No difference was observed between CTL and SMG Exe rats in the DG or SVZ (Figure 6B and C, Bonferroni multiple comparisons test, p > 0.99), showing that 3 weeks of physical exercise was enough to restore survival of newborn cells to a level similar to that of CTL rats. In contrast, a significant difference was observed between CTL Exe and SMG Exe rats in the OB (Figure 6F, Bonferroni multiple comparisons test, p = 0.012), indicating that SMG inhibits the positive effect of physical exercise observed on the survival of the newborn cells in the OB.

No effect of exercise was observed in DCX and NeuN expression in CTL or SMG rats in DG (Figure 6C, three-way ANOVA repeated measures, Exercise effect F(1, 29) = 0.99, p = 0.33). In SVZ, exercise decreased the percentage of DCX cells in both CTL and SMG rats (Figure 6E, three-way ANOVA repeated measures, Exercise effect F(1, 58) = 16.94, p < 0.001; Tukey’s multiple comparisons test, exercise *vs* no exercise, CTL, p = 0.002; SMG, p = 0.002). No difference was observed in NeuN expression (CTL, p > 0.99; SMG, p > 0.99). In the OB, we observed no effect of exercise on DCX and NeuN expression in CTL rats (Figure 6G, three-way ANOVA repeated measures, F(1, 29) = 0,99, p = 0.33, interaction exercise x SMG, F(1, 29) = 5.14, p = 0.03; Tukey’s multiple comparisons test, exercise *vs* no exercise DCX p = 0.98, NeuN p > 0.99). However, exercise decreased the percentage of DCX cells in SMG rats (p = 0.043).

### Neurogenesis related gene expression

#### Simulated microgravity slightly alters the weight gain in rats

In this experiment, dedicated to gene expression analyses, neurogenesis was not assessed. Accordingly, rats were submitted for 7 or 21 days to SMG exposure without EdU/BrdU injections. In these conditions, CTL rats did not lose weight and they continuously gained weight from D1 (Figure 7A, white circle and black arrow; one-sample t-test, D1 to D21 p < 0.05). In contrast, SMG rats lost weight (Figure 7A, red circle) and it took them 8 days to return to their starting weight (Figure 7A, red arrow; one-sample t-test, D1 to D7 p < 0.05; D8 p = 0.2) and 12 more days to increase their weight (one-sample t-test, D9 to D19 p > 0.05, D20 to D21 p < 0.05). A significant difference between CTL and SMG was observed during the 8 first day (two-way repeated measures ANOVA, mixed-effects model, SMG effect F(1, 14) = 88.18, p < 0.001; Bonferroni multiple comparisons test, D1 to D8 p < 0.05; D9 p = 0.08, D10 p = 0.06, D11 p = 0.03, D12 to D21 p > 0.1). The lack of difference between D9 to D21 was mostly due to the low number of animals in both groups (n = 4). A simple linear regression on the 21 days showed that weight gain differed between CTL and SMG rats (F(1, 206) = 10.66, p =0.001). A more detailed analysis showed that this difference was due to the first 2 days of exposure. Indeed, from D3, the simple linear regression showed no difference between CTL and SMG rats (F(1, 174) = 3.47, p = 0.064), indicating that after an initial phase of weight stagnation, SMG rats gain weight in the same way as CTL rats. Physical exercise had no effect on weight gain in CTL rats (Figure 7A, white square; two-way ANOVA, exercise effect F(1, 10) = 4.08, p = 0.07). Simple linear regression showed no difference between rats with or without exercise (F(1, 80) = 1.22, p = 0.27). In SMG rats, physical exercise slightly decreased the weight gain from D3 (Figure 7A, red square; two-way ANOVA, exercise effect F(1, 10) = 9.08, p = 0.013; Bonferroni multiple comparisons test, D1 to D2 p > 0.3, D3 p = 0.043, D4 p = 0.1, D5 p = 0.09, D6 p = 0.005, D7 p = 0.12). However, no difference was observed by simple linear regression between SMG rats with or without exercise (F(1, 73) = 1.88, p = 0.18). These results indicate that physical exercise associated with SMG could impair the rat’s body weight, at least during several days. However, this effect was not due to stress while physical exercise did not change the corticosterone level in rats (data not shown; two-way ANOVA, exercise effect F(1, 12) = 0.05, p = 0.82).

**Figure 7.**
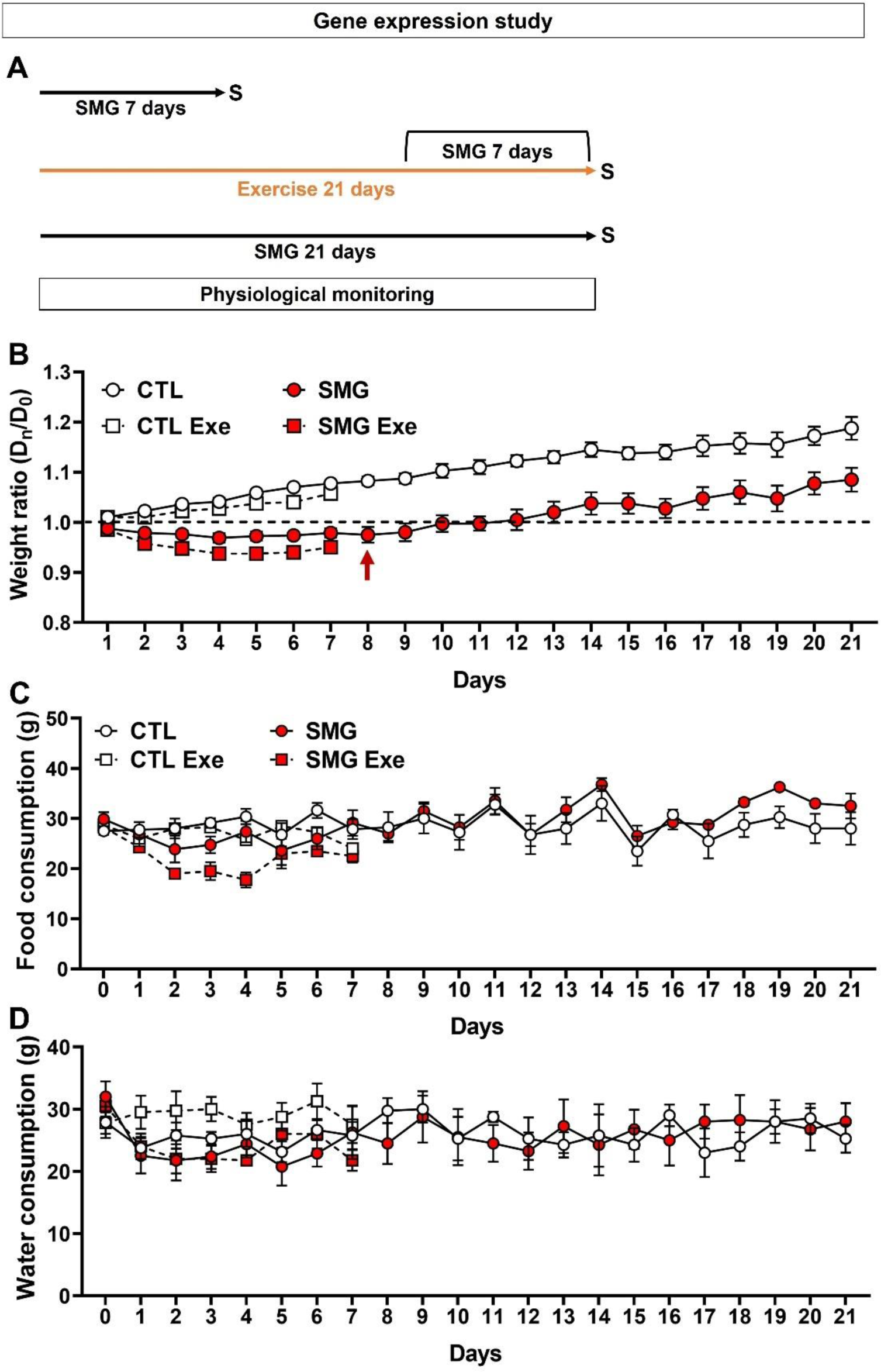
SMG slightly altered the weight gain in rats. **A.** Experimental design. Rats were exposed to SMG during 7 or 21 days. Some rats were exposed to physical exercise during 3 weeks. After 2 weeks, rats were exposed to SMG for 7 days. The physiological parameters of the animals were monitored daily. **B.** Weight of the CTL (white) and SMG (red) rats exposed (square) or not (circle) to physical exercise presented as a ratio compared with the weight at D0. The red arrow marks the point where the SMG rats return to their starting weight. **C.** Food consumption of the CTL (white) and SMG (red) rats exposed (square) or not (circle) to physical exercise. **D.** Water consumption of the CTL (white) and SMG (red) rats exposed (square) or not (circle) to physical exercise. All data are presented as mean ± SEM.

Food and water consumption were stable throughout the experiment in CTL and SMG rats (Food: Figure 7C, two-way ANOVA, mixed-effects model, time effect F(3.16, 27.35) = 3.93, p = 0.017; Dunnett multiple comparisons test, D1 to D21 p > 0.05 in both group; Water: Figure 7D, two-way ANOVA, mixed-effects model, time effect F(2.02, 17.48) = 2.17, p = 0.14). No difference was observed between CTL and SMG rats (Food: Figure 7C, two-way ANOVA, mixed-effects model, SMG effect F(1, 14) = 0.15, p = 0.7; Water: Figure 7D, two-way ANOVA, mixed-effects model, SMG effect F(1, 14) = 0.03, p = 0.86). Physical exercise did not change rats’ food (Figure 7C; three-way ANOVA, exercise effect F(1, 20) = 3.05, p = 0.1) or water consumption (Figure 7D; three-way ANOVA, exercise effect F(1, 20) = 0.63, p = 0.44).

No difference was observed in CTL and SMG rats in temperature (data not shown; two-way ANOVA, mixed-effects model, SMG effect F(1, 14) = 0.51, p = 0.49) or glycemia (data not shown; two-way ANOVA, mixed-effects model, SMG effect F(1, 14) = 0.05, p = 0.83). As previously observed, physical exercise increased the body temperature (data not shown; three-way ANOVA, exercise effect F(1, 20) = 37.77, p < 0.001) and the glycemia (data not shown; D0, two-way ANOVA, exercise effect F(1, 20) = 9.41, p = 0.006).

#### SMG modulates the expression of genes involved in neurogenic processes

The regulation of neurogenesis was first investigated by RT-qPCR using a manufactured plate testing 82 genes known to be related to neurogenesis. After 7 days of SMG exposure, the hippocampal expression of 25 genes were decreased with a 2.5-fold change threshold (Figure 8A and Table 3). After 21 days of SMG exposure, the expression of only 3 genes was increased (Figure 8B and Table 3). The comparison between 7 and 21 days of SMG exposure indicate that the genes *Clcx1* and *Neurog1*, were regulated in an opposite way: 7 days of exposure decreased their expression while 21 days of exposure increased it. These results suggest that SMG had a transient effect on gene expression involved in neurogenesis in the hippocampus.

**Figure 8.**
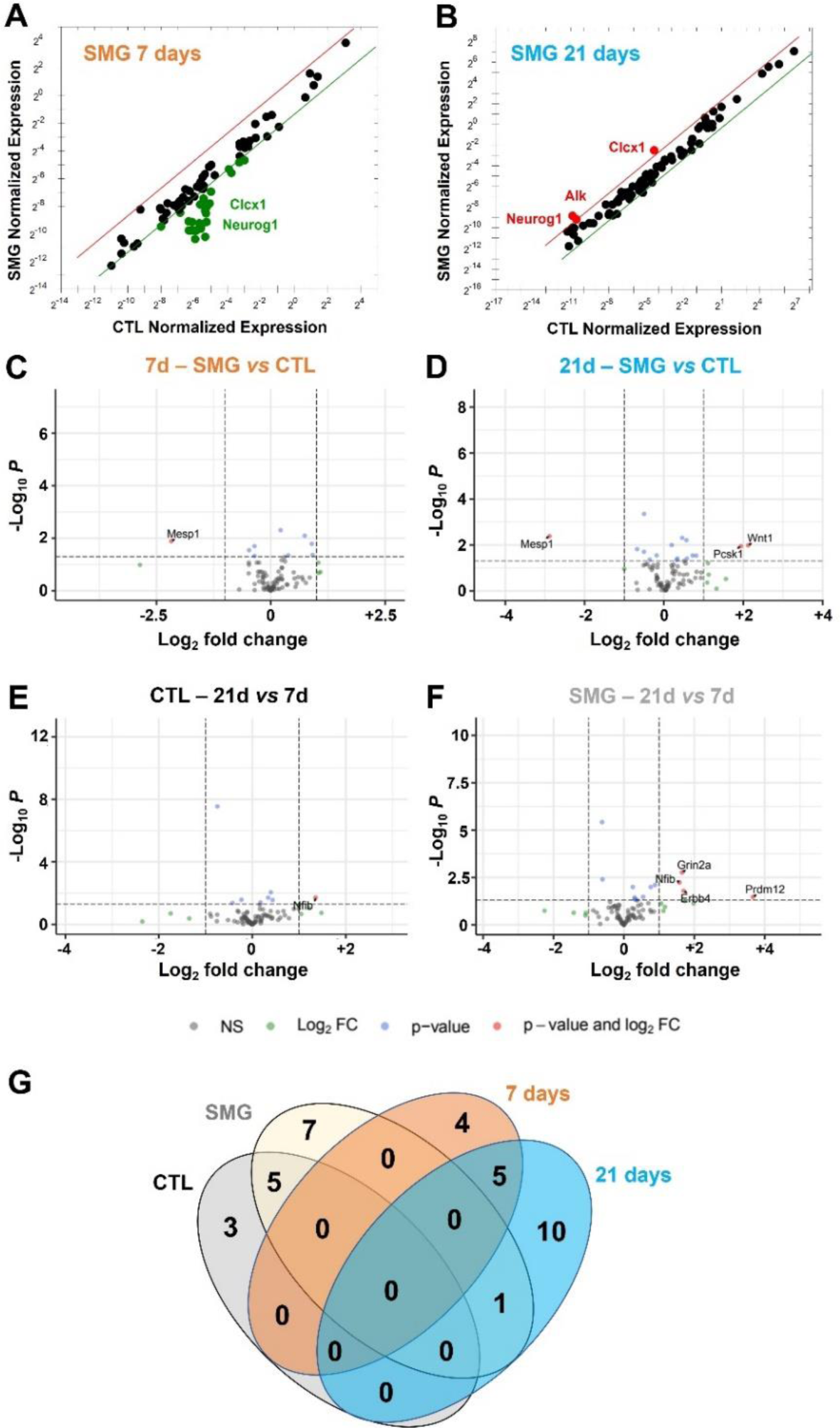
SMG modulates neurogenesis related gene expression in hippocampus. **A-B.** Comparison of normalized expression of genes of rats exposed to SMG during 7 and 21 days, respectively and revealed by RTqPCR. Comparisons are made with control rats. Dots represent the tested genes with a fold change threshold = 2.5 (red dots on the left of the red line are upregulated and green dots on the right green line are downregulated. Graphs are generated by gene study analyses with CFX software (Bio-Rad). **C-D.** Volcano plots of expressions of genes of rats exposed to SMG during 7 or 21 days, respectively, revealed by RNAseq. **E.** Volcano plot of expressions of genes of CTL rats isolated 21 days compared with a 7-day isolation. **F.** Volcano plot of expression of genes of rats exposed to SMG during 21 days compared with a 7-day exposure. In grey (NS) is the group of the non-affected genes, in blue (p < 0.05) the expression of gene is significantly affected, in green the fold is change more than twice (|log2FC| > 2) and finally in red are indicated the gene expression significantly and highly changed (p < 0.05 and |log2FC| > 2). **G.** Venn’s diagram to segregate affected gene populations according to the experimental conditions.

**Table 1.**
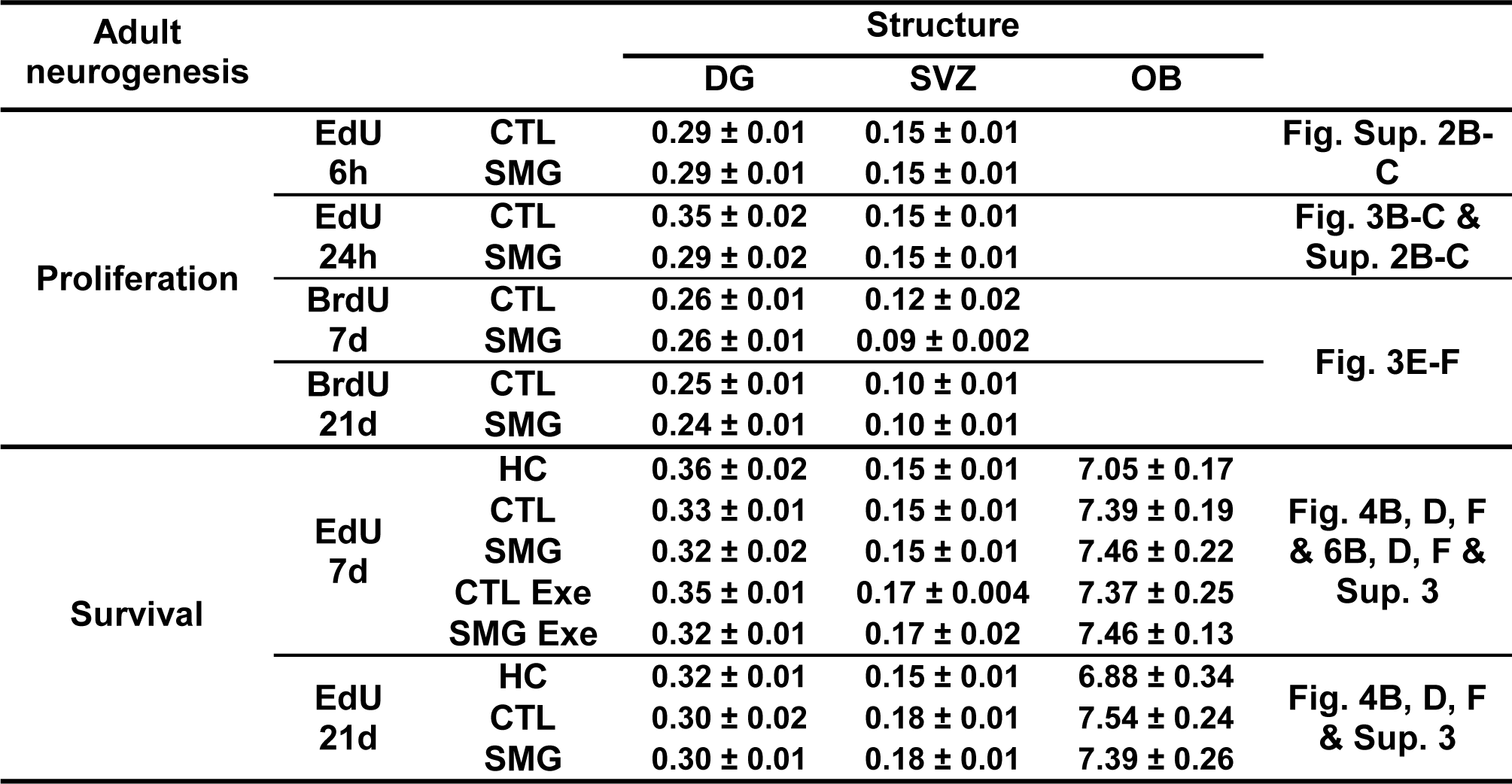
Surface (mm²) analyzed in DG, SVZ and OB for the study of EdU+ and BrdU+ nuclei quantification.

**Table 2.**
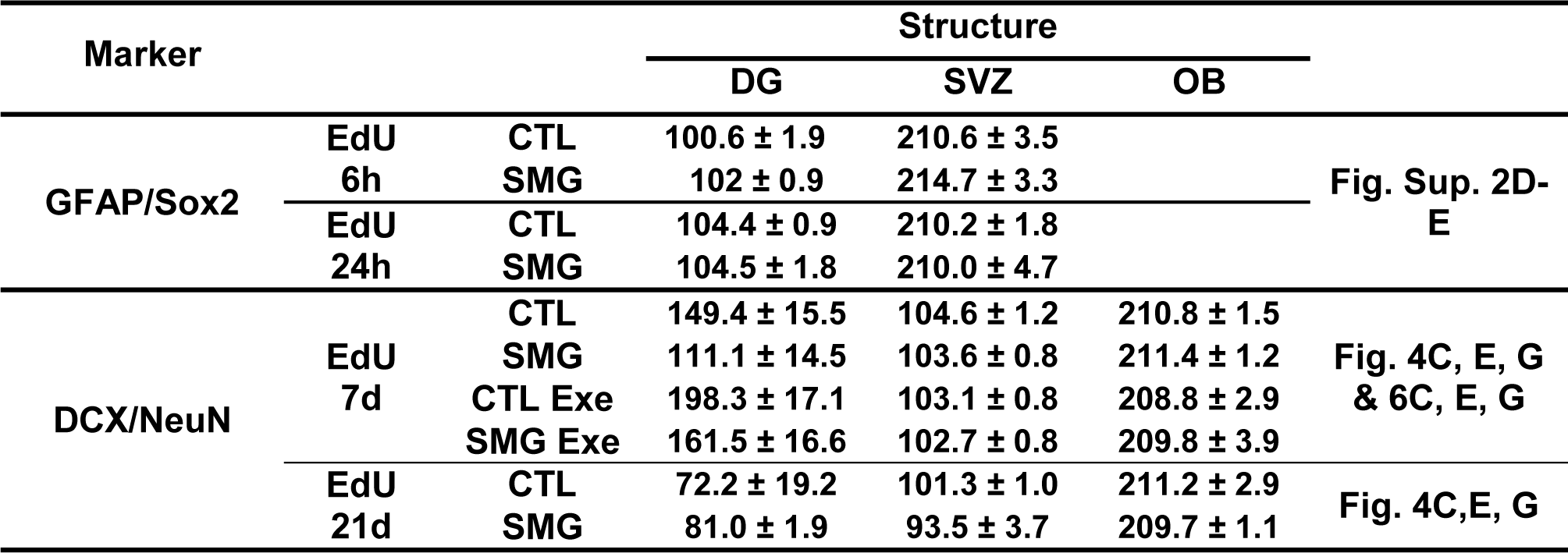
Number of EdU+ cells analyzed in DG, SVZ and OB for the study of GFAP+, Sox2+, DCX+ and NeuN+ cells quantification Marker.

| Marker |  |  | Structure |  |  |  |
| --- | --- | --- | --- | --- | --- | --- |
|  |  |  | DG | SVZ | OB |  |
| GFAP/Sox2 | EdU 6h | CTL | 100.6 ± 1.9 | 210.6 ± 3.5 |  | Fig. Sup. 2D-E |
|  |  | SMG | 102 ± 0.9 | 214.7 ± 3.3 |  |  |
|  | EdU 24h | CTL | 104.4 ± 0.9 | 210.2 ± 1.8 |  |  |
|  |  | SMG | 104.5 ± 1.8 | 210.0 ± 4.7 |  |  |
| DCX/NeuN | EdU 7d | CTL | 149.4 ± 15.5 | 104.6 ± 1.2 | 210.8 ± 1.5 | Fig. 4C, E, G & 6C, E, G |
|  |  | SMG | 111.1 ± 14.5 | 103.6 ± 0.8 | 211.4 ± 1.2 |  |
|  |  | CTL Exe | 198.3 ± 17.1 | 103.1 ± 0.8 | 208.8 ± 2.9 |  |
|  |  | SMG Exe | 161.5 ± 16.6 | 102.7 ± 0.8 | 209.8 ± 3.9 |  |
|  | EdU 21d | CTL | 72.2 ± 19.2 | 101.3 ± 1.0 | 211.2 ± 2.9 | Fig. 4C,E, G |
|  |  | SMG | 81.0 ± 1.9 | 93.5 ± 3.7 | 209.7 ± 1.1 |  |

**Table 3.**
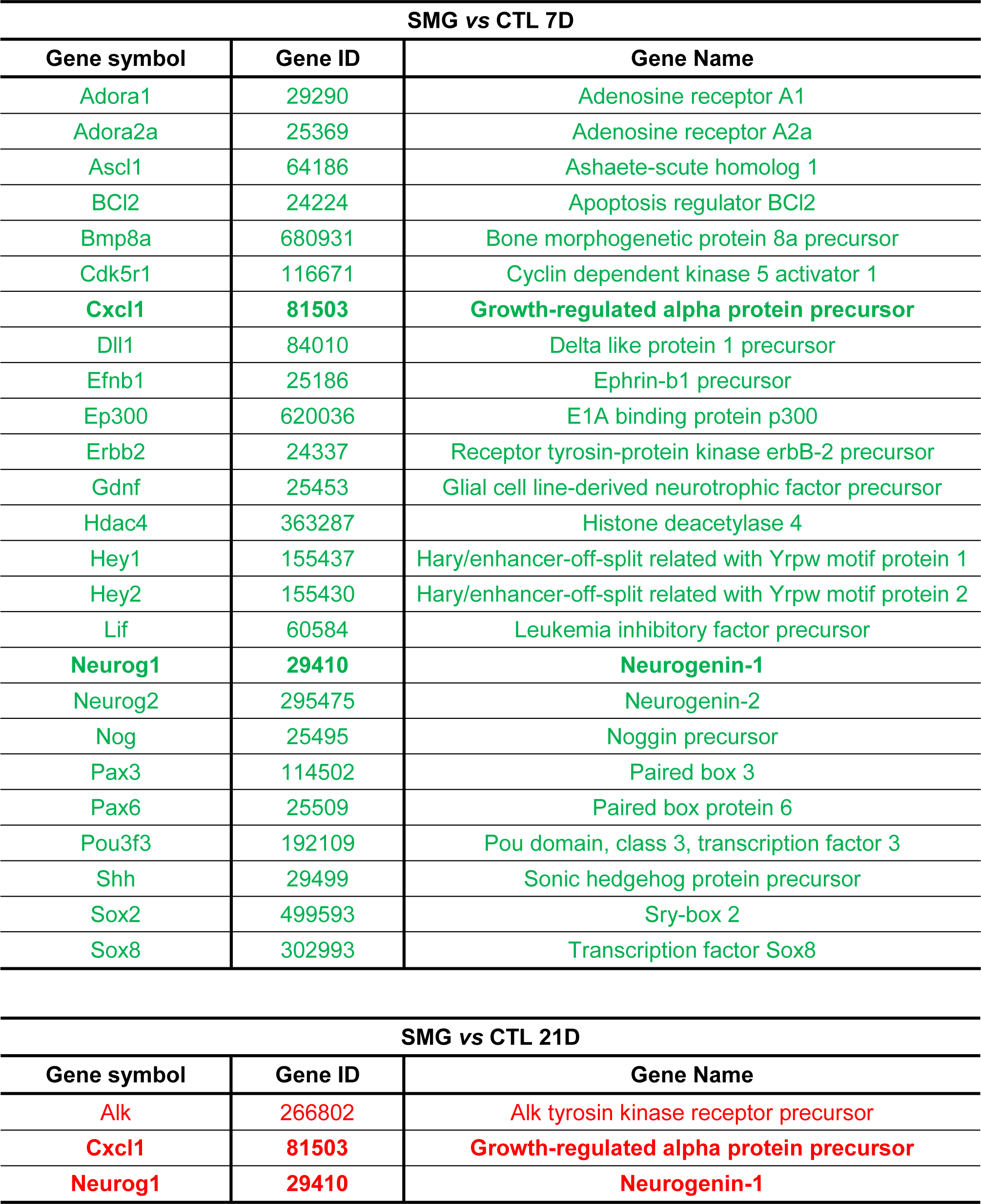
List of genes affected by 7-or 21-day SMG exposure compared to CTL groups by RTqPCR analysis. Down-regulated genes are in green. Up-regulated genes are in red.

To complete the analysis of gene expression implicated in adult neurogenesis, we extracted the genes included in the Go-terms “neurogenesis” containing 89 genes from a large RNA-Seq transcriptomic analysis. We analyzed the transcriptomes in 4 rats per experimental groups and compared the results pairwise. We focused only on the genes whose expression was significantly modified (p < 0.05) and they were highlighted in the volcano plots when the fold change (FC) was |log2FC| > 2). In SMG rats, the expression of 20 genes involved in neurogenesis were significantly modified (SMG *vs* CTL; Fig. 8C-D and Table 4): 4 were affected after 7 days of SMG exposure (*Ascl1, Celsr1, Prdm8* and *Xrcc5*); 11 were affected after the 21-day SMG exposure period (*Adgrv1, Ercc6, Lft20, Nav1, Nav2, Ndufs2, Pcsk1, Tacc1, Trak1* and *Wnt1*); and 5 were affected identically by both delays of exposure (*Fabp7, Mesp1, Nfix, Smarca4, Tuba1a*). Only *Mesp1* was highly decreased for both SMG durations (|log2FC| > 2). We next compared gene expression as a function of the duration of the SMG exposure. In CTL rats, 8 genes were significantly affected by the extensive duration (CTL 21 *vs* 7 days; *Celsr3, Ephb1, Lpar1, Nfib, Ulk4, Erbb3, Marcks* and *Slc1a1*; Fig. 8E and Table 4). The expression of 5 of these genes was also altered in SMG rats (Table 4; *Celsr3, Ephb1, Lpar1, Nfib, Ulk4*), indicating that the altered expression of these genes is not specific to the simulated microgravity condition but could be related to potential side effects of longer isolation in individual cages. Notably, the duration of SMG exposure specifically increased the expression of 7 genes (SMG 21 *vs* 7 days; *Btbd1, Erbb4, Grin2a, Myd88, Nfia, Prdm12, Psen1*; Fig. 8F and Table 4). However, only *Prdm12* expression was highly increased in SMG rats when the SMG period was extended to 21 days (|log2FC| > 2). Results are summarized in the Venn’s diagram (Fig. 8G).

**Table 4.**
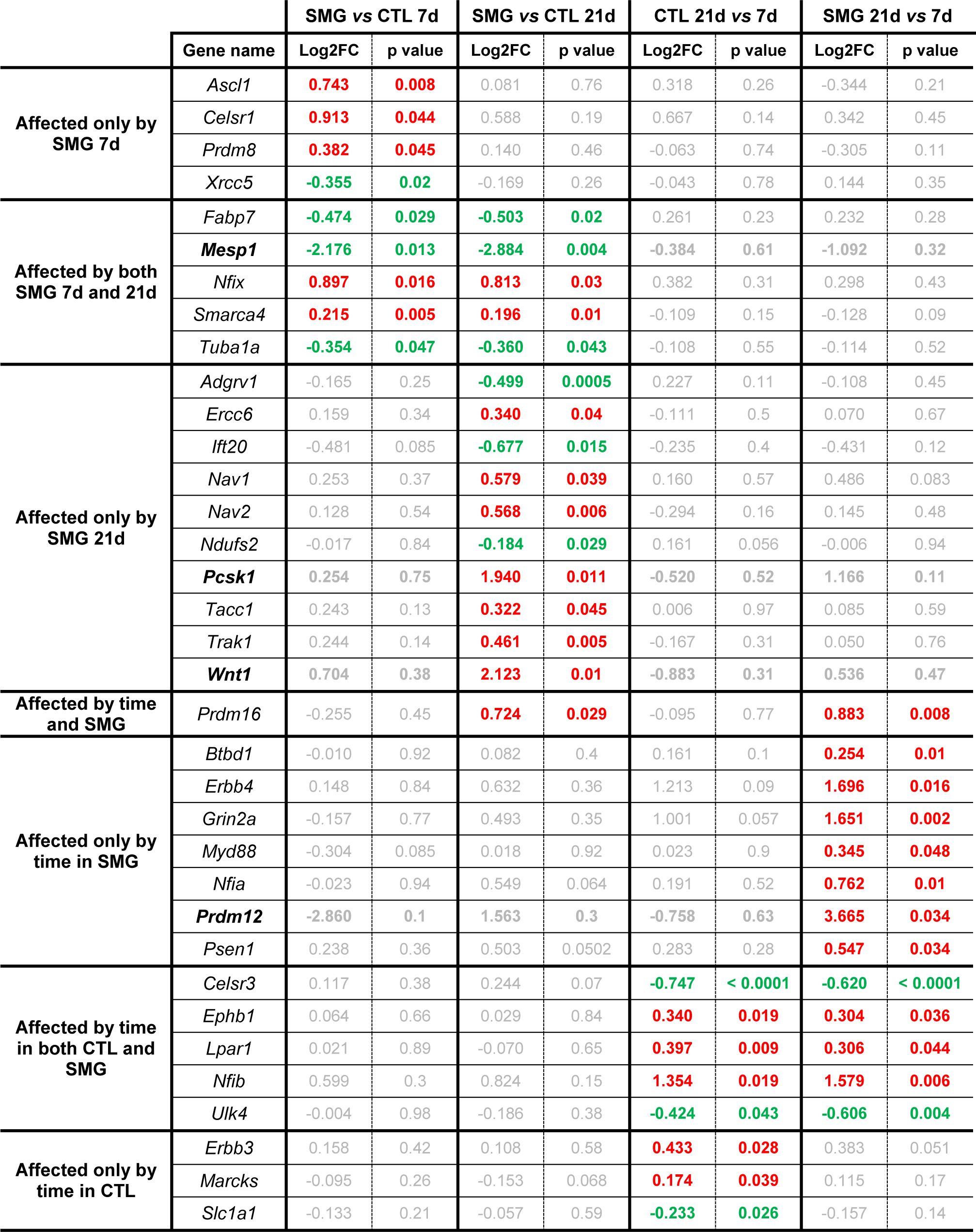
SMG and time effects on the expression of genes included in the neurogenesis GO term list revealed by RNAseq. Down-regulated genes are in green. Up-regulated genes are in red. Insignificant changes are in grey.

|  | Gene name | SMG vs CTL 7d |  | SMG vs CTL 21d |  | CTL 21d vs 7d |  | SMG 21d vs 7d |  |
| --- | --- | --- | --- | --- | --- | --- | --- | --- | --- |
|  |  | Log2FC | p value | Log2FC | p value | Log2FC | p value | Log2FC | p value |
| Affected only by SMG 7d | *Ascl1* | 0.743 | 0.008 | 0.081 | 0.76 | 0.318 | 0.26 | -0.344 | 0.21 |
|  | *Celsr1* | 0.913 | 0.044 | 0.588 | 0.19 | 0.667 | 0.14 | 0.342 | 0.45 |
|  | *Prdm8* | 0.382 | 0.045 | 0.140 | 0.46 | -0.063 | 0.74 | -0.305 | 0.11 |
|  | *Xrcc5* | -0.355 | 0.02 | -0.169 | 0.26 | -0.043 | 0.78 | 0.144 | 0.35 |
| Affected by both SMG 7d and 21d | *Fabp7* | -0.474 | 0.029 | -0.503 | 0.02 | 0.261 | 0.23 | 0.232 | 0.28 |
|  | *Mesp1* | -2.176 | 0.013 | -2.884 | 0.004 | -0.384 | 0.61 | -1.092 | 0.32 |
|  | *Nfix* | 0.897 | 0.016 | 0.813 | 0.03 | 0.382 | 0.31 | 0.298 | 0.43 |
|  | *Smarca4* | 0.215 | 0.005 | 0.196 | 0.01 | -0.109 | 0.15 | -0.128 | 0.09 |
|  | *Tuba1a* | -0.354 | 0.047 | -0.360 | 0.043 | -0.108 | 0.55 | -0.114 | 0.52 |
| Affected only by SMG 21d | *Adgrv1* | -0.165 | 0.25 | -0.499 | 0.0005 | 0.227 | 0.11 | -0.108 | 0.45 |
|  | *Ercc6* | 0.159 | 0.34 | 0.340 | 0.04 | -0.111 | 0.5 | 0.070 | 0.67 |
|  | *Ift20* | -0.481 | 0.085 | -0.677 | 0.015 | -0.235 | 0.4 | -0.431 | 0.12 |
|  | *Nav1* | 0.253 | 0.37 | 0.579 | 0.039 | 0.160 | 0.57 | 0.486 | 0.083 |
|  | *Nav2* | 0.128 | 0.54 | 0.568 | 0.006 | -0.294 | 0.16 | 0.145 | 0.48 |
|  | *Ndufs2* | -0.017 | 0.84 | -0.184 | 0.029 | 0.161 | 0.056 | -0.006 | 0.94 |
|  | *Pcsk1* | 0.254 | 0.75 | 1.940 | 0.011 | -0.520 | 0.52 | 1.166 | 0.11 |
|  | *Tacc1* | 0.243 | 0.13 | 0.322 | 0.045 | 0.006 | 0.97 | 0.085 | 0.59 |
|  | *Trak1* | 0.244 | 0.14 | 0.461 | 0.005 | -0.167 | 0.31 | 0.050 | 0.76 |
|  | *Wnt1* | 0.704 | 0.38 | 2.123 | 0.01 | -0.883 | 0.31 | 0.536 | 0.47 |
| Affected by time and SMG | *Prdm16* | -0.255 | 0.45 | 0.724 | 0.029 | -0.095 | 0.77 | 0.883 | 0.008 |
| Affected only by time in SMG | *Btbd1* | -0.010 | 0.92 | 0.082 | 0.4 | 0.161 | 0.1 | 0.254 | 0.01 |
|  | *ErbB4* | 0.148 | 0.84 | 0.632 | 0.36 | 1.213 | 0.09 | 1.696 | 0.016 |
|  | *Grin2a* | -0.157 | 0.77 | 0.493 | 0.35 | 1.001 | 0.057 | 1.651 | 0.002 |
|  | *Myd88* | -0.304 | 0.085 | 0.018 | 0.92 | 0.023 | 0.9 | 0.345 | 0.048 |
|  | *Nfia* | -0.023 | 0.94 | 0.549 | 0.064 | 0.191 | 0.52 | 0.762 | 0.01 |
|  | *Prdm12* | -2.860 | 0.1 | 1.563 | 0.3 | -0.758 | 0.63 | 3.665 | 0.034 |
|  | *Psen1* | 0.238 | 0.36 | 0.503 | 0.0502 | 0.283 | 0.28 | 0.547 | 0.034 |
| Affected by time in both CTL and SMG | *Celsr3* | 0.117 | 0.38 | 0.244 | 0.07 | -0.747 | < 0.0001 | -0.620 | < 0.0001 |
|  | *Ephb1* | 0.064 | 0.66 | 0.029 | 0.84 | 0.340 | 0.019 | 0.304 | 0.036 |
|  | *Lpar1* | 0.021 | 0.89 | -0.070 | 0.65 | 0.397 | 0.009 | 0.306 | 0.044 |
|  | *Nfib* | 0.599 | 0.3 | 0.824 | 0.15 | 1.354 | 0.019 | 1.579 | 0.006 |
|  | *Ulk4* | -0.004 | 0.98 | -0.186 | 0.38 | -0.424 | 0.043 | -0.606 | 0.004 |
| Affected only by time in CTL | *ErbB3* | 0.158 | 0.42 | 0.108 | 0.58 | 0.433 | 0.028 | 0.383 | 0.051 |
|  | *Marcks* | -0.095 | 0.26 | -0.153 | 0.068 | 0.174 | 0.039 | 0.115 | 0.17 |
|  | *Slc1a1* | -0.133 | 0.21 | -0.057 | 0.59 | -0.233 | 0.026 | -0.157 | 0.14 |

#### Effects of exercise on the neurogenesis related gene expression in the hippocampus

RTqPCR analyses revealed that physical exercise decreased the expression of 27 genes involved in neurogenesis in the hippocampus of CTL rats (Figure 9A and Table 5). In SMG rats exposed to physical exercise, 39 genes showed downregulated expression (Figure 9B and Table 5). Physical exercise induced a lower expression of *ApoE*, *Ll3*, *Pou4f1* and *Egf* in the hippocampus of SMG rats exposed to exercise than in rats exposed only to SMG (figure 9C). In comparison with the control condition, the Venn’s diagrams (Figure 9D) showed that 22/27 genes were exclusively regulated by physical exercise; 24/28 genes were affected by SMG independently of exposure to the exercise countermeasure. These results suggest that exercise and SMG regulated differentially the genes implicated in neurogenesis, and that exercise was not able to reverse the SMG effect on gene expression. In addition, 4 genes (*Cdk5rap2*, *Hdac4*, *Grin1* and *Gpi*) were downregulated similarly by SMG and physical exercise; the gene *Mdk* was less expressed by physical exercise independently of SMG exposure; and 4 genes were regulated by exercise only in rats exposed to SMG (*Egf*, *Creb1*, *Pou4f1* and *Ll3*). Gene populations can be segregated by the modulation of their expression. Accordingly, physical exercise decreased the expression of *Mdk* and *ApoE* in CTL and SMG groups, exacerbated the decrease of *Cxcl1* expression, increased in CTL and decrease in SMG the expression of the genes *Egf*, *Ll3* and *Pouf4f1*, and finally had no effect or only reversed partially the effect of SMG for the others.

**Figure 9.**
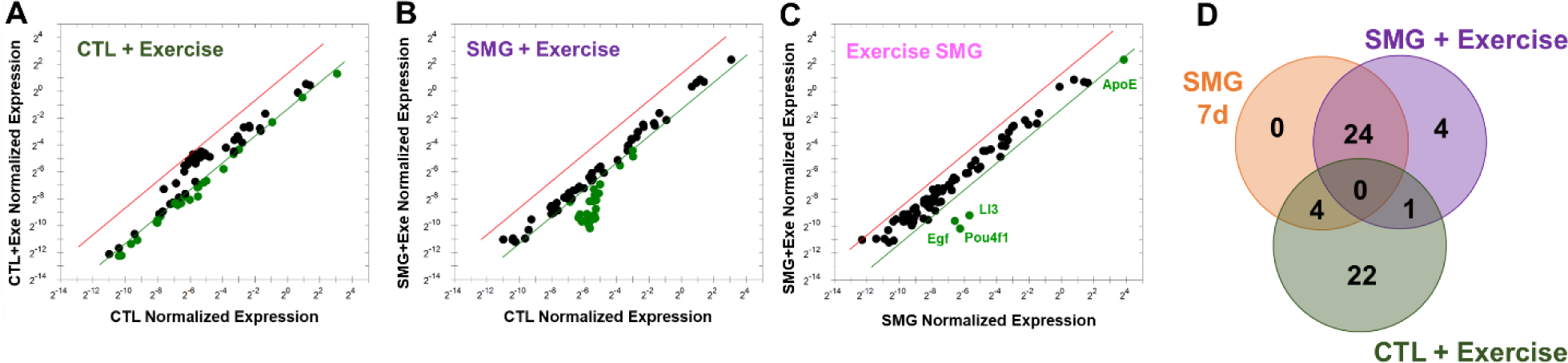
Physical exercise modulates neurogenesis related gene expression in hippocampus by RTqPCR. **A-B.** Comparison of normalized expression of genes in CTL and in SMG conditions, respectively. Comparisons are made with control rats. **C.** Comparison of normalized expression of genes of rats exposed to exercise in SMG conditions. Comparisons are made with SMG rats not exposed to exercise. Dots represent the tested genes with a fold change threshold = 2.5 (green dots on the right green line are downregulated). Graphs are generated by gene study analyses with CFX software (Bio-Rad). **D.** Venn’s diagram to segregate gene populations affected by experimental conditions.

**Table 5.**
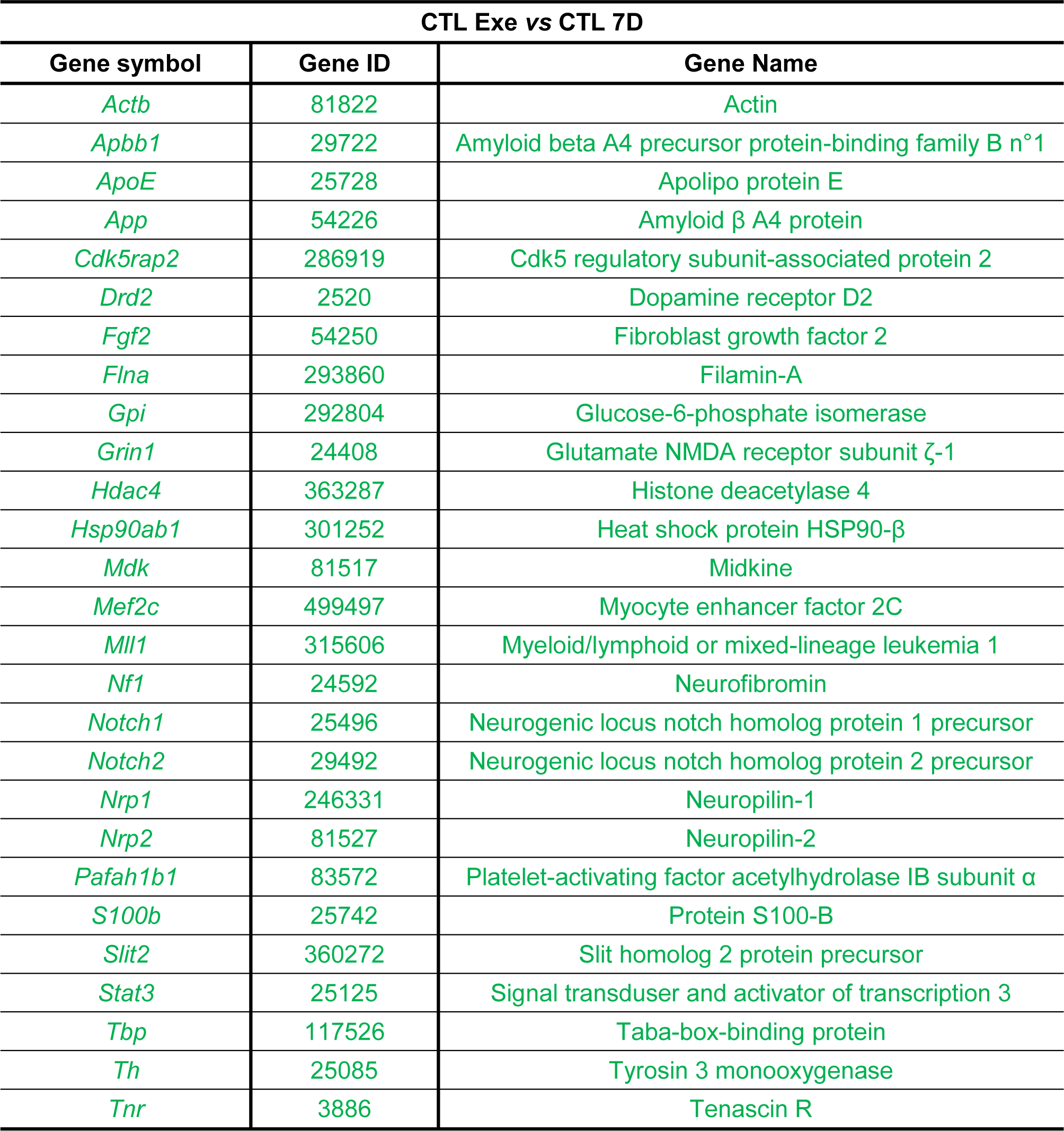
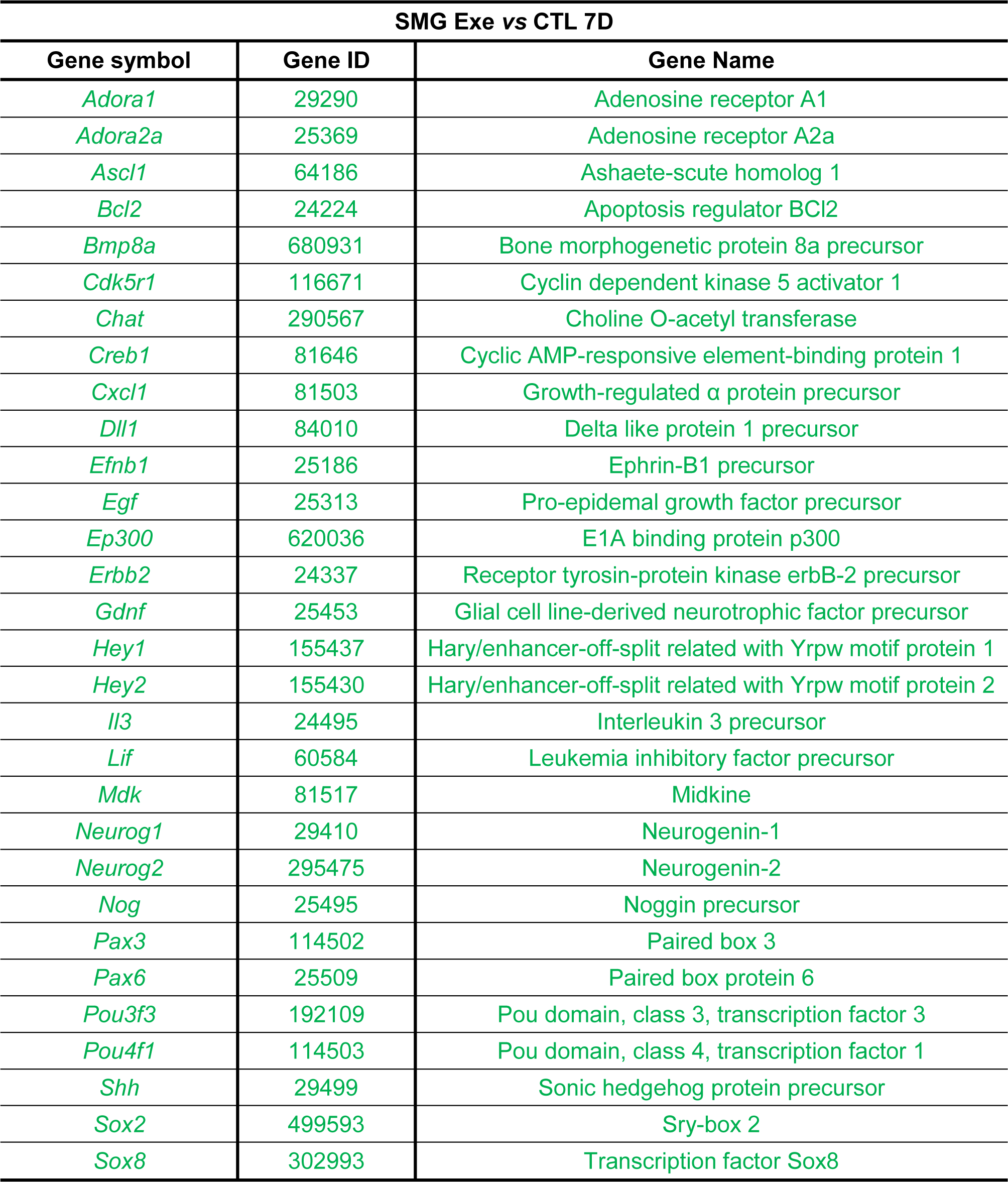
List of genes affected by physical exercise in control and rats exposed to SMG during 7 days by RTqPCR analysis. Down-regulated genes are in green.

| CTL Exe vs CTL 7D |  |  |
| --- | --- | --- |
| Gene symbol | Gene ID | Gene Name |
| *Actb* | 81822 | Actin |
| *Apbb1* | 29722 | Amyloid beta A4 precursor protein-binding family B n°1 |
| *ApoE* | 25728 | Apolipo protein E |
| *App* | 54226 | Amyloid β A4 protein |
| *Cdk5rap2* | 286919 | Cdk5 regulatory subunit-associated protein 2 |
| *Drd2* | 2520 | Dopamine receptor D2 |
| *Fgf2* | 54250 | Fibroblast growth factor 2 |
| *Flna* | 293860 | Filamin-A |
| *Gpi* | 292804 | Glucose-6-phosphate isomerase |
| *Grin1* | 24408 | Glutamate NMDA receptor subunit ζ-1 |
| *Hdac4* | 363287 | Histone deacetylase 4 |
| *Hsp90ab1* | 301252 | Heat shock protein HSP90-β |
| *Mdk* | 81517 | Midkine |
| *Mef2c* | 499497 | Myocyte enhancer factor 2C |
| *Mll1* | 315606 | Myeloid/lymphoid or mixed-lineage leukemia 1 |
| *Nf1* | 24592 | Neurofibromin |
| *Notch1* | 25496 | Neurogenic locus notch homolog protein 1 precursor |
| *Notch2* | 29492 | Neurogenic locus notch homolog protein 2 precursor |
| *Nrp1* | 246331 | Neuropilin-1 |
| *Nrp2* | 81527 | Neuropilin-2 |
| *Pafah1b1* | 83572 | Platelet-activating factor acetylhydrolase IB subunit α |
| *S100b* | 25742 | Protein S100-B |
| *Slit2* | 360272 | Slit homolog 2 protein precursor |
| *Stat3* | 25125 | Signal transducer and activator of transcription 3 |
| *Tbp* | 117526 | Taba-box-binding protein |
| *Th* | 25085 | Tyrosin 3 monooxygenase |
| *Tnr* | 3886 | Tenascin R |

| SMG Exe vs CTL 7D |  |  |
| --- | --- | --- |
| Gene symbol | Gene ID | Gene Name |
| *Adora1* | 29290 | Adenosine receptor A1 |
| *Adora2a* | 25369 | Adenosine receptor A2a |
| *Ascl1* | 64186 | Ashaete-scute homolog 1 |
| *Bcl2* | 24224 | Apoptosis regulator BCL2 |
| *Bmp8a* | 680931 | Bone morphogenetic protein 8a precursor |
| *Cdk5r1* | 116671 | Cyclin dependent kinase 5 activator 1 |
| *Chat* | 290567 | Choline O-acetyl transferase |
| *Creb1* | 81646 | Cyclic AMP-responsive element-binding protein 1 |
| *Cxcl1* | 81503 | Growth-regulated $\alpha$ protein precursor |
| *Dll1* | 84010 | Delta like protein 1 precursor |
| *Efnb1* | 25186 | Ephrin-B1 precursor |
| *Egf* | 25313 | Pro-epidermal growth factor precursor |
| *Ep300* | 620036 | E1A binding protein p300 |
| *ErbB2* | 24337 | Receptor tyrosin-protein kinase erbB-2 precursor |
| *Gdnf* | 25453 | Glial cell line-derived neurotrophic factor precursor |
| *Hey1* | 155437 | Hary/enhancer-off-split related with Yrpw motif protein 1 |
| *Hey2* | 155430 | Hary/enhancer-off-split related with Yrpw motif protein 2 |
| *Il3* | 24495 | Interleukin 3 precursor |
| *Lif* | 60584 | Leukemia inhibitory factor precursor |
| *Mdk* | 81517 | Midkine |
| *Neurog1* | 29410 | Neurogenin-1 |
| *Neurog2* | 295475 | Neurogenin-2 |
| *Nog* | 25495 | Noggin precursor |
| *Pax3* | 114502 | Paired box 3 |
| *Pax6* | 25509 | Paired box protein 6 |
| *Pou3f3* | 192109 | Pou domain, class 3, transcription factor 3 |
| *Pou4f1* | 114503 | Pou domain, class 4, transcription factor 1 |
| *Shh* | 29499 | Sonic hedgehog protein precursor |
| *Sox2* | 499593 | Sry-box 2 |
| *Sox8* | 302993 | Transcription factor Sox8 |

**Table 6.** Exercise and SMG effects on the expression of genes included in the neurogenesis GO term list revealed by RNAseq. Down-regulated genes are in green. Up-regulated genes are in red. Insignificant changes are in grey.

|  | Gene name | SMG vs CTL 7d |  | CTL + Exe vs CTL 7d |  | SMG + Exe vs CTL 7d |  | SMG + Exe vs SMG 7d |  | SMG + Exe vs CTL + Exe |  |
| --- | --- | --- | --- | --- | --- | --- | --- | --- | --- | --- | --- |
|  |  | Log2FC | p value | Log2FC | p value | Log2FC | p value | Log2FC | p value | Log2FC | p value |
| Affected by exercise in CTL | *Adgrv1* | -0.165 | 0.25 | -0.308 | 0.032 | -0.215 | 0.13 | -0.050 | 0.73 | 0.093 | 0.52 |
| Affected by exercise in both | *Dagla* | 0.297 | 0.052 | 0.399 | 0.009 | 0.355 | 0.02 | 0.058 | 0.71 | -0.044 | 0.78 |
|  | *Grin2a* | -0.157 | 0.77 | 1.186 | 0.024 | 1.125 | 0.032 | 1.282 | 0.015 | -0.061 | 0.91 |
| Affected only by SMG without exercise | *Celsr1* | 0.913 | 0.044 | 0.158 | 0.73 | 0.305 | 0.5 | -0.608 | 0.18 | 0.147 | 0.75 |
|  | *Mesp1* | -2.176 | 0.013 | -1.071 | 0.17 | -0.927 | 0.23 | 1.249 | 0.16 | 0.144 | 0.86 |
|  | *Nfix* | 0.897 | 0.017 | 0.520 | 0.17 | 0.575 | 0.13 | -0.323 | 0.39 | 0.054 | 0.88 |
|  | *Prdm8* | 0.382 | 0.045 | 0.277 | 0.15 | 0.248 | 0.19 | -0.134 | 0.48 | -0.030 | 0.88 |
|  | *Smarca4* | 0.215 | 0.005 | 0.079 | 0.3 | 0.120 | 0.12 | -0.095 | 0.21 | 0.041 | 0.59 |
|  | *Xrcc5* | -0.355 | 0.02 | -0.013 | 0.93 | -0.290 | 0.056 | 0.066 | 0.67 | -0.276 | 0.068 |
| Affected by SMG and exercise in SMG condition | *Tuba1a* | -0.354 | 0.047 | -0.135 | 0.45 | -0.447 | 0.012 | -0.092 | 0.61 | -0.312 | 0.08 |
|  | *Fabp7* | -0.474 | 0.029 | -0.220 | 0.3096 | -0.535 | 0.013 | -0.060 | 0.78 | -0.315 | 0.15 |
|  | *Ascl1* | 0.743 | 0.008 | -0.084 | 0.77 | 0.133 | 0.64 | -0.610 | 0.027 | 0.217 | 0.45 |
| Affected by exercise in SMG | *Brinp1* | 0.170 | 0.19 | 0.209 | 0.1 | 0.335 | 0.009 | 0.164 | 0.19 | 0.125 | 0.33 |
|  | *Marcks* | -0.095 | 0.26 | -0.118 | 0.16 | -0.180 | 0.034 | -0.085 | 0.32 | -0.061 | 0.47 |
|  | *Fat4* | 1.041 | 0.085 | 1.101 | 0.068 | 1.237 | 0.04 | 0.195 | 0.74 | 0.136 | 0.82 |

RNAseq analyses revealed that the expression of *Adgrv1* was significantly decreased by physical exercise only in CTL rats (Table 5). The expression of *Dagla* and *Grin2a* was increased by physical exercise in both SMG and CTL rats but only *Grin2a* expression was also increased by exercise in comparison with SMG rats not exposed to exercise (Fig. 10C and Table 5).

**Figure 10.**
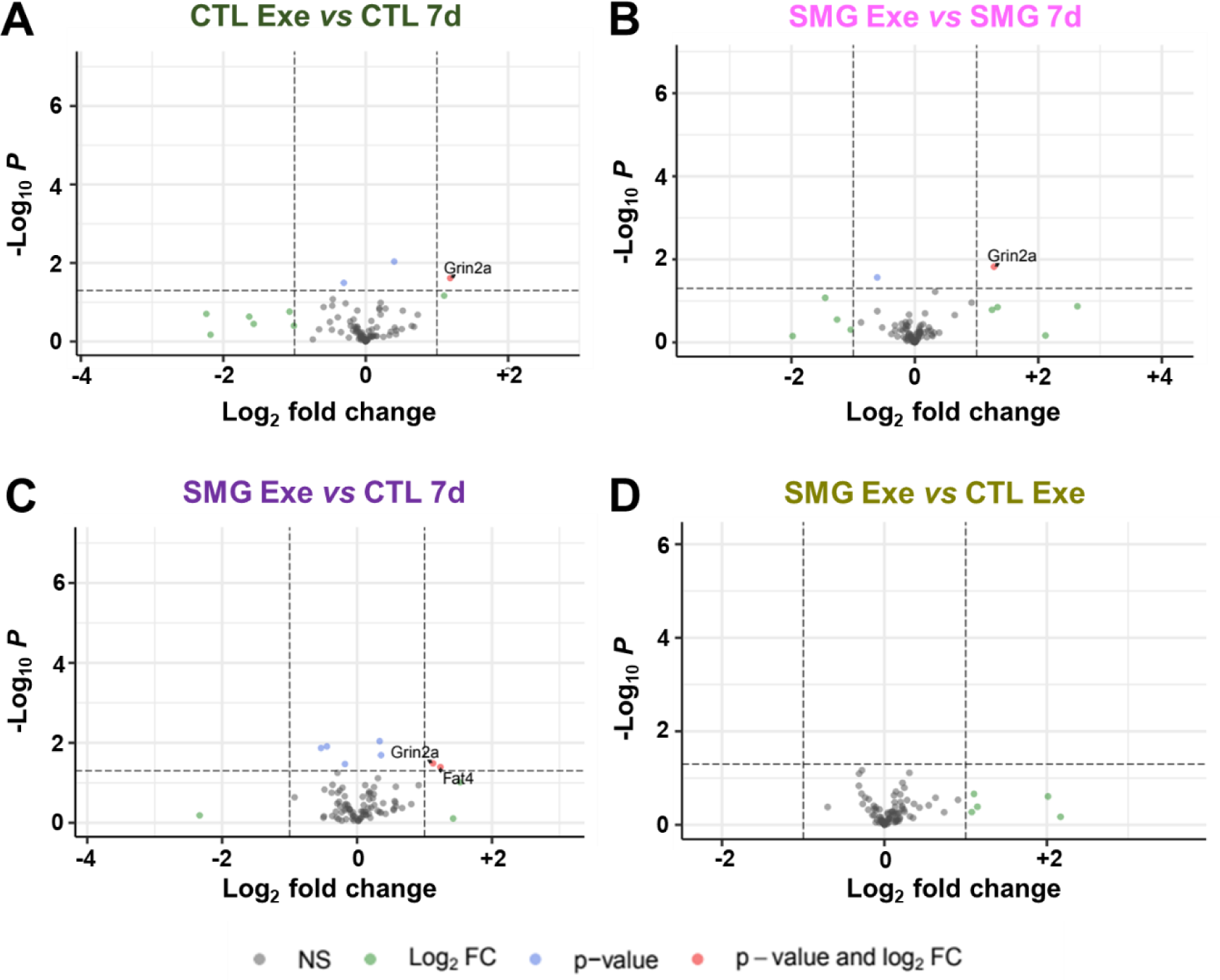
Physical exercise modulates neurogenesis related gene expression in hippocampus by RNAseq. **A-B.** Volcano plots of expression of genes of rats exposed to exercise in CTL **(A)** or SMG **(B)** conditions. **C-D.** Volcano plot of expression of neurogenesis related genes in hippocampus of rats exposed to exercise in SMG conditions compared with CTL rats without **(C)** or with **(D)** exercise. In grey (NS) is the group of the non-affected genes, in blue (p < 0.05) the expression of gene is significantly affected, in green the fold is change more than twice (|LnFC| > 2) and finally in red are indicted the gene expression significantly and highly changed (p < à.05 and |LnFC| > 2).

While 7 days of SMG exposure decreased the expression of 4 genes involved in neurogenesis (Fig. 8C and Table 4-5; *Mesp1, Xrcc5*, *Tuba1a* and *Fabp7*) and increased the expression of 5 other genes (Fig. 8C and Table 4-5; *Ascl1, Celsr1, Prdm8, Nfix* and *Smarca4*), the expression of *Tuba1a* and *Fabp7* was similarly decreased in SMG rats performing exercise (Table 5). While the expression of Ascl1 was decreased in the SMG condition, physical exercise reversed its expression (Table 5). Finally, the expression of *Brinp1*, *Marcks* and *Fat4* was changed in SMG rats submitted to exercise compared to the CTL condition.

## Discussion

The present study found significant reduction of newborn cell proliferation in DG and newborn cell survival in DG and SVZ after 7 days of exposure to SMG in male rats. This neurogenesis impairment was only transient as it was no longer observable after a SMG period of 21 days. Physical exercise used as a countermeasure was able to restore adult neurogenesis in the DG of rats exposed to SMG.

### Physiological parameters are impacted by simulated microgravity in the HU model

It is well-known that microgravity impacts the soleus muscle in astronauts during spaceflights and in animal models of SMG (Ohira, 2000). Reduction in muscle mass observed in the present study and is in accordance with several previous investigations that have found muscle atrophy after 7 days (Bederman et al., 2015; Ulanova et al., 2019; Yoshihara et al., 2021), 2 weeks (Chowdhury et al., 2016, 2013; Momken et al., 2011; Mortreux et al., 2021; Ohira et al., 1992; Yasuhara et al., 2007) or one month of hindlimb suspension (Tahimic et al., 2019). Physical exercise administered for two weeks in an activity wheel can counteract this deleterious effect (Yasuhara et al., 2007). Concurring with this, we also observed an increase in the soleus muscle weight of our rats exercising for 3-week on a running treadmill. We did not observe any difference in the distance covered by control and SMG rats during our imposed treadmill sessions. However, in spontaneous locomotor activity tests during which animals exposed to SMG can freely access an activity wheel, detailed analyses of locomotion patterns showed multiple long-term deficits, such as a shorter distance covered, a lower number of movements and their duration, and a reduced vertical activity (Yasuhara et al., 2007). These alterations were observed immediately upon the end of SMG exposure and persisted 2 weeks later, indicating that the effects of SMG on locomotion are long-lasting (Yasuhara et al., 2007).

Simulated microgravity in the HU model leads to a reduction in weight gain of rats. This alteration in body weight during suspension has been documented on several previous occasions in rodents (Bederman et al., 2015; Bo et al., 2019; Chowdhury et al., 2013, 2016; Feng et al., 2016a; Momken et al., 2011; Mortreux et al., 2021; Tahimic et al., 2019; Tanaka et al., 2013; Tsvirkun et al., 2012; Ulanova et al., 2019; Wang et al., 2021; Yasuhara et al., 2007). As previously shown (Chowdhury et al., 2013, 2016; Feng et al., 2016a; Mortreux et al., 2021; Tsvirkun et al., 2012), the consumption of water and food in our study was not impacted by simulated microgravity during hindlimb suspension. It is interesting to note that, despite identical food and water consumption, rodents subjected to SMG show lower weight gain at least at the start of the procedure. This suggests either metabolic changes (Feng et al., 2016b), or an increase in energy expenditure (Lew et al., 2009; Momken et al., 2011) in rats undergoing SMG. In our experiments, EdU injections led to a transient reduction in food and water consumption, associated with reduced weight gain in both CTL and SMG rats. Since these injections represent a significant intake of body fluid, it may explain the drop in water consumption the day after the injections. In addition, injections could generate a stress, known to induce hypophagia (Vallès et al., 2000).

### Simulated microgravity impairs adult neurogenesis

We observed that SMG impaired newborn cell proliferation in the DG by 40% following a 7-day exposure period to HU but not after a shorter period of 6 or 24 hours. It has been previously observed that SMG decreased by 50% the newborn cell proliferation in DG after 2 weeks exposure in the HU model in rats (Yasuhara et al., 2007), by two third after 2 weeks exposure in the HU model in mice (Adami et al., 2018) or by 40% after 6 weeks exposure in the head-down bed rest model in rhesus macaque (Zhang et al., 2019). On the contrary, 3-day exposure to SMG using the HU model did not decrease the number of Ki67^+^ cells in the DG (Berezovskaya et al., 2021). These results indicate that it does require some time of exposure to microgravity before effects on the proliferation of new cells can be observed. Nevertheless, in our study, no effect was observed on newborn cell proliferation after 21 days of exposure to SMG. In addition, we observed that the levels of newborn cell proliferation were not similar in CTL rats between 7- and 21-days of SMG exposure (although the difference (p < 0.12) failed to reach significance). This lack of effect after 21 days of exposure to SMG could be masked by the reduced number of newborn cells produced in CTL rats. This reduction in newborn cell proliferation in CTL rats may be due to social isolation, which can alter the proliferation of the newborn cells in the hippocampus (Holmes, 2016; Lu et al., 2003).

We also observed that SMG reduced newborn cell survival in DG by a third and in SVZ by 20% after 7-day exposure to HU. However, no effect of SMG on adult neurogenesis was observed after 21 days of exposure to HU, indicating a transient effect. Exposure to SMG in HU model during 3 days induced a decrease in number of immature newborn neurons in the DG in rats (Berezovskaya et al., 2021). In rhesus macaques, it has been observed a 50% decrease of newborn cell survival in the DG after 6 weeks exposure to SMG in the head-down bed rest model (Zhang et al., 2019). In this last study, monkeys were severely restrained on the bed, which can induce stress associated with increase in plasma hormones involved in response to stress, known to reduce adult hippocampal neurogenesis (Schoenfeld and Gould, 2012). Moreover, in humans, the plasma level of cortisol was increased in volunteers during bed rest experiments (Choukèr et al., 2001). In our HU model, any major stress effects on adult neurogenesis appear unlikely because we did not observe an increase of the corticosterone level in rats submitted to SMG. In addition, the effect of SMG on newborn cell survival observed in HU rats was not due to social isolation, as no difference was observed between isolated and group-housed rats (Fig. Sup. 3). Finally, no effect of SMG was observed in the OB. This is consistent with previous study showing no difference on the density of newborn neurons in OB of female mice after a 13-day spaceflight (Latchney et al., 2014).

### Molecular mechanisms underlying the effect of microgravity on adult neurogenesis

The effect of SMG on adult neurogenesis seems to be attributable to modifications in biological and molecular properties of neural stem cells. This cell type derived from the SVZ of mice exposed to HU model during 2 weeks showed reduced proliferation capability, altered cell cycle and incomplete differentiation/maturation (Adami et al., 2018). However, 3-day exposure to SMG in the HU model is not enough to induce deregulation of the expression of Sox2 (Berezovskaya et al., 2021) which controls proper self-renewal of neural stem cells (Episkopou, 2005). In addition, HU-induced SMG induces a decrease in the cerebral expression of several factors known to play a role in adult neurogenesis regulation such as NR2A/NR2B NMDA receptors (Berezovskaya et al., 2021; Xiang et al., 2019; Zhai et al., 2019), insulin-like growth factor-1 (IGF-1) (Mysoet et al., 2014), brain derived neurotrophic factor (BDNF) (Bo et al., 2019; Wu et al., 2017; Zhai et al., 2020) or p-CREB (cAMP-response element binding protein) (Wu et al., 2017; Xiang et al., 2019).

The results of our RTqPCR and transcriptomic analyses suggested that the effects of the exposure to 7 and 21 days to hindlimb suspension affected differently the neurogenesis related gene expression. After 7 days of SMG, some genes implicated in neurogenesis were partially downregulated, suggesting a global impairment of neurogenesis pathways. But after 21 days of SMG, the expression of these genes was upregulated. This result suggests that during the first week of SMG, the adult neurogenesis process is globally impacted by simulated microgravity, but after this initial period, some compensatory mechanisms were engaged to restore the original level of hippocampal integration of newborn neurons. In comparison, after 28 days of hindlimb suspension, proteomic analysis suggested that the expression of few (close to 10%) proteins involved in neurogenesis were affected (Ji et al., 2022).

In our analysis, we observed that simulated microgravity regulated several genes known to be involved in neurogenesis. It has been shown that these genes, as *EdbB4, Ascl1, Pax6, EphB, Cxcl1, Bcl2, Tuba1a or Nfix,* are notably involved in the organization of the neurogenic niche (Ghashghaei et al., 2006), in neural stem cells quiescence (Andersen et al., 2014; Blomfield et al., 2019) and maintenance (Curto et al., 2014; Liu et al., 2016; Sakamoto et al., 2003), in neural stem cell self-renewal (Bauer and Patterson, 2006; Palma et al., 2005), in progenitor cell proliferation (Amador-Arjona et al., 2011; Cabello-Rivera et al., 2019; Chumley et al., 2007; Huang et al., 2018; Kim et al., 2005; Palma et al., 2005) and differentiation (Curto et al., 2014), in neuroblast maturation and survival (Harris et al., 2018; Kuhn et al., 2005), in neurite growth (Shima et al., 2007), in neuronal polarization (Christova et al., 2023), in neuronal migration (Chumley et al., 2007; Feng et al., 2012; Keays et al., 2010; Zalucki et al., 2019), or in the integration of newly generated neurons (Bonafina et al., 2019; Curto et al., 2014).

To better understand the regulation of gene expression induced by microgravity, the experiments should be investigated more precisely by single cell RNAseq (Overbey et al., 2022; Willis et al., 2020) and localization experiments based on the result of the bulk analysis reported here. In fact, SMG should have different effects on neuronal lineages depending also on the duration of SMG exposure. Moreover, many other pathways concerning neuron activity should be analyzed in more details, including the role of inflammatory factors, microglia and oligodendrocytes and factors released by endothelium (Lin et al., 2020; Yan et al., 2021).

### Physical exercise as a countermeasure to preserve adult neurogenesis during spaceflight

Physical activity is the most effective way to limit the adverse effects of microgravity on the human body during spaceflight (English et al., 2020; Loehr et al., 2015; Scott et al., 2023), including treadmill training (which provides cardiovascular, muscular and skeletal exercise) (Petersen et al., 2016), and represent an essential part of the astronauts’ daily routine. Physical exercise (wheel or treadmill) is a well-proven neurogenic stimulus in rodents, promoting neuronal progenitor proliferation and newborn cell survival, and influencing the morpho-functional maturation process of newborn neurons (Farmer et al., 2004; Ferreira et al., 2011; Inoue et al., 2015; Jin et al., 2017; Lattanzi et al., 2022; Leasure and Jones, 2008; Lee et al., 2013; Okamoto et al., 2012; Patten et al., 2013; Ra et al., 2002; Trejo et al., 2001; Uda et al., 2006; Yasuhara et al., 2007).

Treadmill speed has been shown to induce different effects on adult hippocampal neurogenesis. At low-or mild-intensity speed, i.e., between 10 and 30 cm/s, several studies have shown a 1.5 to 2-fold increase in the level of newborn cells produced and surviving in the DG (Ferreira et al., 2011; Okamoto et al., 2012; Ra et al., 2002; Trejo et al., 2001; Uda et al., 2006). However, intense-intensity speed, i.e., superior to 40 cm/s, induces an increase of the proliferation of newborn cells but no effect on their long-term survival (Inoue et al., 2015; Ra et al., 2002). Based on these previous results and on the fact that rats would be subjected to exercise during SMG exposure, we chose a mild-intensity speed for our study. In control rats, we reproduced the 1.5 to 2-fold increase of the newborn cells’ survival in the DG (Fig. 6), indicating that our exercise protocol has the expected effect on adult neurogenesis. Moreover, we observed a 1.5-fold increase of the number of newborn neurons in the OB. This result is surprising as exercise has been shown to induce an increase in adult neurogenesis in the hippocampus but not in the OB (Brown et al., 2003). In our study, the animals were isolated, leading to a reduction in social interactions and a depletion of olfactory stimuli. During treadmill sessions, the animals may have left a variety of olfactory marks in a confined space possibly leading to an increase in olfactory stimulation and consequently an increase in adult neurogenesis in the OB. This effect has already been observed in a confined space module left on the ground (Latchney et al., 2014).

The major finding of our study is that physical exercise completely restores newborn cell survival impaired by SMG in the DG and SVZ. A positive effect of physical exercise has already been observed in rats exposed to the HU model, without however completely restoring newborn cell proliferation level in DG or SVZ (Yasuhara et al., 2007). The difference observed between this previous study and ours is probably due to the type of exercise, i.e., activity wheel *vs* treadmill, its intensity, i.e., free access *vs* imposed speed, and duration, i.e., 3 hours/day during 2 weeks *vs* 30 minutes/day during 3 weeks. Moreover, in our experiment, rats underwent exercise sessions before and during SMG exposure, whereas in the study by Yasuhara and colleagues, rats were exposed to physical exercise only after SMG exposure (Yasuhara et al., 2007). In addition, they did not observe a positive effect on adult neurogenesis when SMG exposure was discontinued for 2 weeks without physical exercise (Yasuhara et al., 2007). This indicates that adult neurogenesis remains impacted for a long time after exposure to SMG. Thus, it is therefore unlikely that in our protocol, the observed effect of physical exercise on adult neurogenesis survival in a SMG context is linked to the fact that the animals are no longer suspended for 30 minutes a day for exercise session. Moreover, our results also confirm those obtained with dynamic foot stimulation of the plantar surface in rats during SMG exposure that also prevents the reduction of DCX cells in the DG (Berezovskaya et al., 2021). Altogether, it seems that preventive exposure to physical exercise and/or during SMG exposure are more effective in preventing the impact of SMG on adult neurogenesis than during the recovery phase after SMG exposure.

Molecular mechanisms underlying the effect of physical exercise on adult neurogenesis are related to many factors, such as BDNF (Farmer et al., 2004; Jin et al., 2017), NMDA receptors (Farmer et al., 2004), fibroblast growth factor 2 (FGF-2) (Gómez-Pinilla et al., 1997), vascular endothelial growth factor (VEGF) (Fabel et al., 2003) or IGF-1 (Carro et al., 2000; Trejo et al., 2001). The neurogenic niche in the DG is organized in close association to endothelial cells forming a supportive vascular niche in which VEGF and IGF-1 serve as supporting factors (Carro et al., 2000; Fabel et al., 2003; Trejo et al., 2001). Physical exercise in rats submitted to simulated microgravity in the HU model maintains levels expression of NMDA receptors and VEGF comparable to control rats (Berezovskaya et al., 2021; Yasuhara et al., 2007). In this study, the expression of these genes was not affected by SMG. The transcriptomic analysis indicated that the effects of the exercise did not reverse strictly the effects of SMG on neurogenesis related genes but may have acted *via* modulation of different transduction pathways.

### Limitations and perspectives

This study primarily focused on the effects of SMG on the proliferation and survival of newborn neurons in the adult brain of resting rats. How SMG may affect adult neurogenesis induced by behavioral challenges deserves further investigations. For instance, it will be particularly interesting to explore how SMG may affect the functional integration of new neurons in relation to learning and memory functions in rats exposed to HU and submitted to hippocampal-dependent tasks. It is well known that adult neurogenesis plays an important role in learning and memory processes (Gros et al., 2015), including the formation of enduring memories (Kee et al., 2007; Trouche et al., 2009; Veyrac et al., 2013) or the maintenance and update of remote memory reconsolidation (Lods et al., 2021).

In this study, we limited our transcriptome analysis to adult neurogenesis. It would be relevant to extend the analysis to the entire hippocampal transcriptome and to focus more specifically on the genes involved in learning and memory functions. iTRAQ-based proteomics analyses identified changes in metabotropic and ionotropic glutamate receptor pathway associated with impairment in spatial learning and memory in rats exposed to 28 days of SMG (Wang et al., 2017). Also, after 7 days of hindlimb suspension in mice, proteomic analysis in hippocampus revealed major losses of proteins involved in metabolism or in structural proteins (Sarkar et al., 2006).

To better adapt future physical exercise protocols during spaceflights and long-term missions, the effects of exercise as a countermeasure should be more thoroughly characterized. In particular, it would be interesting to determine whether physical exercise has a differential impact on adult neurogenesis before or during exposure to SMG. Moreover, the effects of spaceflights on physiological parameters depend on the duration of exposure to microgravity. To restore physiological functions altered by spaceflights, the optimal exercise modalities in terms of intensity and duration before and/or during spaceflight, and including individual parameters concerning muscular capacities, should be identified to design the most appropriate exercise protocols.

In addition, it will be important to investigate additional contributing factors likely to alter adult neurogenesis during spaceflights. Combining several stress factors such as radiations (McNerlin et al., 2022) or hypomagnetic field (Zhang et al., 2021) will be needed to better model the environmental conditions experienced by astronauts during space missions. Another major point to consider is the sex effect. Indeed, adult neurogenesis is not regulated in the same manner in male and female rodents and the effects of spaceflights are not exactly similar. RNA-Seq analysis could be a pertinent tool to investigate the cross effects between sex and SMG on adult neurogenesis.

## Materials and methods

### Animals

Adult male Long Evans rats (Charles River, 7 weeks on arrival, n = 124) were use throughout. Rats were initially housed in groups, 2 per cage, according to the calculation of the size of the cage as a function of their weight (model GR900, surface: 904 cm², Techniplast) and were maintained under standard condition in a temperature- and humidity-controlled experimental room. The room was under a 12h light/dark cycle (light onset 7.30AM) and experiments were performed during the light phase of the cycle. Food (A04 product, SAFE) and tap water were available *ad libitum* during the study. The rats were handled once daily to minimize stress and to familiarize them with the male and female experimenters. Enrichment was also used to reduce stress (piece of wood to chew on, wood chips, cardboard tunnel). All experiments were in accordance with the European Communities Council Directive (2010/63/EU Council Directive Decree) regarding the care and use of laboratory animals. The procedures were approved by the Ethical Committee (CCEA-050, project #28854) of the University of Bordeaux. All efforts were made to minimize the number of animals and their suffering, and to maximize their well-being during the experiments.

### Experimental model for simulated microgravity

To simulate microgravity, we used the HU rat model (Imetronic®, Figure 1A), modified from the Morey-Holton method (Globus and Morey-Holton, 2016; Morey-Holton et al., 2005; Morey-Holton and Globus, 2002) and described in our previous study (Gros et al., 2021). Briefly, the tails of the rats were cleaned and protected with medical adhesive tape. A metal ring with a flexible cord was attached to the rat’s tails with tape. The cord was clipped on a pulley attached to a mobile metal bar inserted in the roof of the cages. Rats were able to move freely in a 360° arc in the cage (surface 1050 cm²) with their forelimbs. The hindlimbs were kept above the cage floor. The head-down tilt position was near 40-45°.

### Experimental design

After acclimation to the vivarium conditions (UMR 5293 Institute, University of Bordeaux, agreement A32-063-940) and daily handling for 10 days, the rats were divided into several groups to evaluate adult neurogenesis (n = 97) and gene expression (n = 24) (Figure 1). After an acclimation period in individual cages for 3 days, the hindlimb-suspended rats were maintained under simulated microgravity for 6h to 21 days. For the adult neurogenesis study, rats were exposed to SMG for 6 hours (CTL n = 8, SMG n = 8), 24 hours (CTL n = 8, SMG n = 8), 7 days (HC n = 10, CTL n = 8, SMG n = 8), or 21 days (HC n = 6, CTL n = 6, SMG n = 7) after nucleoside analogue (EdU or BrdU) injections. Physical exercise was used as a countermeasure in rats exposed 7 days to SMG (CTL Exe n = 10, SMG Exe n = 10). For the gene expression study, rats were exposed to SMG for 7d (CTL n = 4, SMG n = 4, CTL Exe n = 4, SMG Exe n = 4) or 21d (CTL n = 4, SMG n = 4). The animals were monitored daily to ensure access to water and food and verify their well-being and that their hindlimbs were not touching the floor grid. The control rats were kept isolated in standard cages with an identical grid to suspended cages to ensure similar sensory inputs between conditions. We did not observe any suffering in the hindlimb-suspended rats. The suspension was stopped for 2 out to 49 rats due to tail swelling during the experiment after 5 and 13 days of hindlimb suspension. Moreover, one rat died before the beginning of the experiment. These 3 rats were excluded from the study.

### Physical exercise (treadmill)

Prior to the experiment, rats (n =28 – 20 rats for the adult neurogenesis study and 8 rats for the gene expression study) were habituated to the treadmill (Imetronic®, Fig. 5A) for 2 days during which they were allowed to explore the treadmill at low speeds (5 to 10 cm/s) for 10 min. The speed of the treadmill was then increased to 20 cm/s for three consecutive daily sessions of 10 minutes. Rats in the exercise groups were placed in a treadmill and underwent forced running for 30 min a day for 3 weeks: 2 weeks prior to the SMG protocol and 1 week during the SMG exposure. During the first week, the speed was set at 25 cm/s. During the second and third weeks, the speed was increased to 30 cm/s. Rats were carefully monitored and the speed of the treadmill was adapted to accommodate the behavior of the animals during the running session, in particular for the rats under SMG, to avoid stress and injuries and to maintain the animal welfare. The speed of the treadmill and/or the duration of the sessions was adapted for 12 rats (4 CTL and 8 SMG – minimum 15 cm/s – minimum 15 min/session), i.e., 42.9% of the rats involved in the physical exercise protocol, for 1 to 6 running sessions. Running experiments take place during the light phase between 8.30 AM and 5.00 PM. Control rats remained in their home cage near the treadmill.

### Physiological measurements

The weight and body temperature of the rats and their food/water consumption was recorded daily. For rats submitted to physical exercise in the treadmill, measures were taken after the exercise session. Glycemia was measured in blood with the Accu-check Performa device (Roche), the day of the isolation, just before HU suspension and just after the end of the suspension procedure 6h, 24h, 7 days or 21 days later. Blood was collected by puncturing the end of the tail of each animal. The corticosterone concentrations were measured by ELISA (K014-Arbor Assay LLC) following the instructions of the supplier, in the supernatant of the blood collected just before perfusion and after centrifugation at 10,000 rpm for 10 min at 4 °C. The supernatant was stored at −20 °C until use. Results were read using a microplate reader (FLUOstar® Omega, BMG Labtech, Champigny-sur-Marne, France) at 450 nm. To check the effect of SMG on muscle weight, two different muscles were collected, the soleus muscle from the hind limbs and the extensor carpi radialis longus muscle from the fore limbs, in CTL (n = 4) and SMG (n = 4) rats. Muscles were weighed directly after collection on the same precision scale.

### Adult neurogenesis study EdU and BrdU administration

To study adult newborn cell proliferation, rats received one intraperitoneal injection of 5-Ethynyl-2’-deoxyuridine (EdU, 200 mg/kg in 0.9% NaCl, BOC Sciences 61135-33-9) and were killed 6 (CTL n = 8, SMG n = 8) or 24h (CTL n = 8, SMG n = 8) later. For adult newborn cell survival, rats were given two EdU injections (200 mg/kg in 0.9% NaCl) at 6h interval on a single day and were killed 7 (HC n = 10, CTL n = 8, SMG n = 8, Exo CTL n = 10, Exo SMG n = 10) or 21 (CTL n = 4, SMG n = 4) days later. These rats also received one intraperitoneal injection of 5-Bromo-2’-deoxyuridine (BrdU, 200 mg/kg in 0.9% NaCl, BOC Sciences 59-14-3) 24h before sacrifice to study adult newborn cell proliferation after several days of simulated microgravity exposure and to reduce the number of animals used for this study.

### Tissue collection and sectioning

Rats were deeply anesthetized with an intraperitoneal injection of mixed Exagon (40 mg/kg) and Lurocaïne (20 mg/kg) solution and then perfused transcardially with a 0.9% NaCl solution during 20 min and then a 4% ice-cold paraformaldehyde solution in 0.1 M phosphate buffer (PB), pH 7.4 during 30 min at 13 ml/min. Brains were removed and post-fixed overnight in the same perfusion solution at 4 °C, immersed for 5 days in PB containing 30% sucrose, frozen in chilled 2-methylbutane at – 50 °C and preserved at - 20 °C. Coronal serial 40-µm-thick free-floating sections were cut with a cryostat (Leica CM3050S) for the olfactory bulb (OB), SVZ and hippocampus area. Brain sections were stored at −20 °C in a cryoprotectant solution until use.

### Immunochemistry

Revelations of EdU+, BrdU+, Sox2+, GFAP+, DCX+ and NeuN+ cells were performed on floating sections. After several rinses in phosphate buffer solution (PBS 0.1 M) to remove the cryoprotection solution, sections were pre-incubated in Target Retrieval Solution (S1700, Dako) for 20 min at 80 °C. After cooling for 10 min, sections were treated with 0.5% Triton in PBS for 20 min. EdU revelation was performed using the Click-it^TM^ EdU Alexa Fluor^TM^ 555 imaging kit (C10338, Invitrogen), incubated for 30 min. For BrdU revelation, two supplementary steps were applied: sections were incubated for 15 min in 2N HCl at room temperature to denature the DNA strands and allow the antigenic sites to become accessible to the antibody, followed by 15 min in 0.1M Boric acid, pH 8.5 to reduced pH to a neutral value. After PBS washes, sections were incubated for 60 min in 10% goat serum (CAECHV00-0U, EuroBio), 0.2% Triton (Triton X-100, 9002-93-1, Sigma), and 0.1% BSA (04-100-812-C, Euromedex) to block nonspecific binding, and incubated overnight at room temperature in a rabbit anti-BrdU primary antibody (1:250, GTX128091, GeneTex), and/or a chicken anti-GFAP (1:4000, Ab4674, Abcam), a rabbit anti-Sox2 (1:250, GTX101507, GeneTex), a guinea-pig anti-DCX (1:2000 for OB, 1:500 for SVZ and hippocampus, Abcam ab2253), a rabbit anti-NeuN (1:2000, GTX132974, GeneTex). Sections were then incubated for 2h at room temperature in a goat anti-rabbit secondary Cy3 antibody (1:250, A120-201C3, Bethyl), and/or goat anti-chicken Cy5 (1:500, A30-206C5, Bethyl), goat anti-rabbit 488 (1:500, A11008, Invitrogen-ThermoFisher), a goat anti-guinea-pig 488 (1:500 for OB, 1:250 for SVZ and hippocampus, A11073, Invitrogen-ThermoFisher), a goat anti-rabbit Cy5 (1:500, A120-201C5, Bethyl). Sections were rinsed, incubated 10 min with Hoechst® (1:2000) in PBS, rinsed in PB, mounted and coverslipped in Fluoromount (FP-48331, Interchim).

### Image acquisition, quantification and analysis

All cell counts were conducted by an experimenter blind to the experimental conditions.

#### EdU+ and BrdU+ nuclei quantification

Labeled profiles in hippocampus were counted with a Leica microscope (DM6000 B) coupled with a JAI camera (Apex Series AT-140GE) and a mapping software (Mercator Pro, Explora Nova). The surface area of the DG was traced at objectives 2.5X (NA 0.07) and 10X (HC PL AP0 NA 0.4) and reported in Table 1. Counting of EdU+ and BrdU+ cells was done in the DG of the dorsal hippocampus, from Bregma - 2.52 mm to −4.56 mm, using a mean of 7 sections per animal spaced by 240 µm, at the objective 20X (PL Ap0 NA 0.6) and 40X (PL AP0 NA 0.75). The density of EdU+ of BrdU+ cells per mm² in the DG was obtained by dividing the number of EdU+ or BrdU+ cells detected by the surface counted.

Labeled profiles in OB and SVZ were counted with a slide scanner (Nanozoomer 2.0 HT, Hamamatsu) at objective 20X. The surface area of the SVZ and OB was traced using the mapping QuPath software and reported in Table 1. EdU+ and BrdU+ nuclei were automatically counted with QuPath (v0.2.3, 2020 QuPath developers, University of Edinburgh) in the OB using a mean of 10 sections and in the SVZ using a mean of 9 sections per animal spaced by 240 µm. The TRITC fluorescence was automatically detected based on a fixed threshold apply for all the animal to avoid bias. For SVZ, the fluorescence was strong and clustered in large areas of several EdU+ or BrdU+ cells. Based on this observation, the detected fluorescence surface was divided by the area of an SVZ cell (41 µm²) to obtain the number of EdU+ or BrdU+ cells. Then the number of EdU+ or BrdU+ cells was divided by the SVZ surface counted to obtain the density of EdU+ or BrdU+ cells per mm².

#### GFAP+, Sox2+, DCX+ and NeuN+ cells quantification

Sections were analyzed using a spinning disk microscope (Leica DMI8, Leica Microsystems, Wetzlar, Germany) equipped with a confocal Scanner Unit CSU-W1 T2 (Yokogawa Electric Corporation, Tokyo, Japan). Images were acquired using an 40X (HCX PL Ap0 NA 1.25) objective and a sCMOS Prime 95B camera (Photometrics, Tuscon, USA). The LASER diodes used were at 405 nm (100 mW), 488 nm (400 mW), 561 nm (400 mW) and 642 nm (100 mW). Z-stacks were done with a galvanometric stage (Leica Microsystems, Wetzlar, Germany). The system was controlled by MetaMorph software (Molecular Devices, Sunnyvale, USA).

### Statistical analysis

All data were averaged across animals within each experimental condition and are presented as mean ± SEM. The normality of the data was checked using Shapiro-Wilk test. All statistical analysis were performed with GraphPad Prism 9 (version 9.3.1). Statistical significance was set at p < 0.05.

Muscle weight was analyzed using unpaired t-test. The effect of physical exercise on muscle weight was analyzed using two-way ANOVA followed by Tukey test for multiple comparisons. Corticosterone concentration was analyzed using two-way ANOVA to compare CTL and SMG or CTL Exe and SMG Exe groups, and using one-way ANOVA to analyze the time effect in each group. The weight of the rats was analyzed as a ratio (*i.e.,* their weight at D_n_ compared with their weight at the start of the experiment, D0). The evolution of the body weight over time was analyzed using one-sample t-test against the value 1. The difference between CTL and SMG groups was analyzed using two-way ANOVA followed by post-hoc tests for multiple comparisons. The effect of physical exercise was analyzed using three-way ANOVA. Food and water consumption, temperature and glycemia were analyzed using two-way ANOVA. The evolution of the consumption over time was analyzed using Dunnett test for multiple comparisons in relation to D0. The difference between CTL and SMG groups was analyzed using Bonferroni test for multiple comparisons. The effect of physical exercise was analyzed using three-way ANOVA followed by Tukey test for multiple comparisons.

Adult neurogenesis was analyzed using unpaired t-test (or Mann-Whitney test for non-normal data) or using two-way ANOVA followed by Bonferroni test for multiple comparisons. Percentage of EdU/BrdU+ cells expressing GFAP+, Sox2+, DCX+ and NeuN+ was analyzed using paired or unpaired t-test or two-or three-way ANOVA with repeated measures followed by Bonferroni test for multiple comparisons.

### Gene expression study Tissue collection

At the end of the SMG exposure (7d, CTL n = 4, SMG n = 4, CTL Exo n = 4, SMG Exo n = 4; 21d, CTL n = 4, SMG n = 4), rats were deeply anesthetized with an intraperitoneal injection of mixed Exagon (40 mg/kg) and Lurocaïne (20 mg/kg) solution and then perfused transcardially with a 0.9% NaCl solution during 10 min at 13 ml/min. Brains were collected and hippocampi were dissected and placed in tubes containing tri-reagent (TR118, Molecular Center, Inc.). Samples were preserved at - 20°C until use.

### Preparation of RNA samples for transcriptomic assays

Hippocampus samples were dissected homogenized with BeadBug6 (3 cycles, speed 4350, time 60s, Benchmak distributed by Dominique Dutscher). Isolation of the total RNA from each hippocampus was performed with Direct-zol RNA miniprep (ZR2053, Zymo) following the supplier procedures. The concentration of RNA was measured by spectrophotometry (NanoDrop 2000, ThermoFisher) for each animal. RNA quality was checked using Lab Chip GX touch HT (PerKin Elmer) and RNA chip (Xmark – High sensitivity). We harmonized the concentration of all sample (∼ 300 ng/µl) and mix the same volume of samples per condition to obtain one RNA solution for each experimental condition.

### Quantitative real-time Polymerase Chain reaction after transcription (RT-QPCR)

To determine the expression of genes involving in adult neurogenesis, we used the Neurogenesis (SAB Target List) R96 Predesigned 96-well panel (10046994, Bio-Rad). We performed the reverse transcription (RT) reaction with 200 ng/µl of total RNA with the iScript gDNA clear advanced kit (Bio-Rad 172-5035) and the real time qPCR was performed with the Sso Advanced SYBR Green® Supermix (Bio-Rad 172-5270) in the CFX96^TM^ Real-Time System thermocycler (Bio-Rad). All samples were analyzed in triplicates and plate preparations were performed randomly with the same lot of PCR mix. Analyses were performed using CFX Maestro (Bio-Rad). Gene expression was normalized by reference gene (Hprt1), the most stable gene in the context of our study. A change in gene expression was considered only for a modification higher than 2.5-fold compared with the control group.

### Bulk RNA-sequencing

cDNA libraries were synthesized using 500 ng of RNA from each sample with the « Illumina Stranded mRNA Prep Ligation » kit according to the supplier instructions. Briefly, the first step consisted on capturing mRNAs with magnetic beads targeting their poly-A tail. Then, mRNAs were fragmented, reverse transcribed into double strand cDNA and 3’ adenylated adapters and anchors were ligated to both ends. Finally, the cDNA library was amplified by PCR with a number of PCR cycles adapted to the start material quantity. Libraries quality was checked with Xmark chip on the Lab Chip GX touch HT (Perkin Elmer). After quantification of individual libraries by q-PCR using the NEBNext library quant kit (New England BioLabs), all the samples were pooled in an equimolar manner. The quality and quantity of the pool were checked by qPCR (RocheLC480) and Lab Chip GX touch HT (Perkin Elmer) before sequencing. mRNA sequencing was performed using Nextseq 2000 Illumina (paired-end 2×100 bp) with a minimum of 30 million reads per sample in the PGTB facility (INRAE - Pierroton).

### Bioinformatics

Bulk-RNA sequencing reads were first pre-processed to ensure data quality. Thus, fastq files quality was analyzed using the fastp software with default parameters (Chen, 2023) and only reads longer than 20bp were kept. Then, reads were aligned to the rat genome (Rnor_6.0 genome assembly) using the splice-aware mapper STAR with default parameters (Dobin et al., 2013) and gene counts were summarized using the GenCounts options of STAR. To determine differentially expressed genes, we used the DESeq2 algorithm in R with default parameters (Love et al., 2014). Gene expression data were displayed as volcano plots using the EnhancedVolcano package in R (https://github.com/kevinblighe/EnhancedVolcano.).

## Acknowledgements

Part of the experiments (sequencing of the RNA-seq experiment) were performed at the PGTB (doi:10.15454/1.5572396583599417E12) with the help of Erwan Guichoux and Zoé Delporte. We thank the staff of the PIVE-EXPE animal facility for animal care. Microscopy was done in the Bordeaux Imaging Center a service unit of the CNRS-INSERM and Bordeaux University, member of the national infrastructure France BioImaging supported by the French National Research Agency (ANR-10-INBS-04). The help of Magalie Mondin is acknowledged and we thank Mélodie Ambroset for her assistance on image analysis.

---

**Figure Sup. 1.**
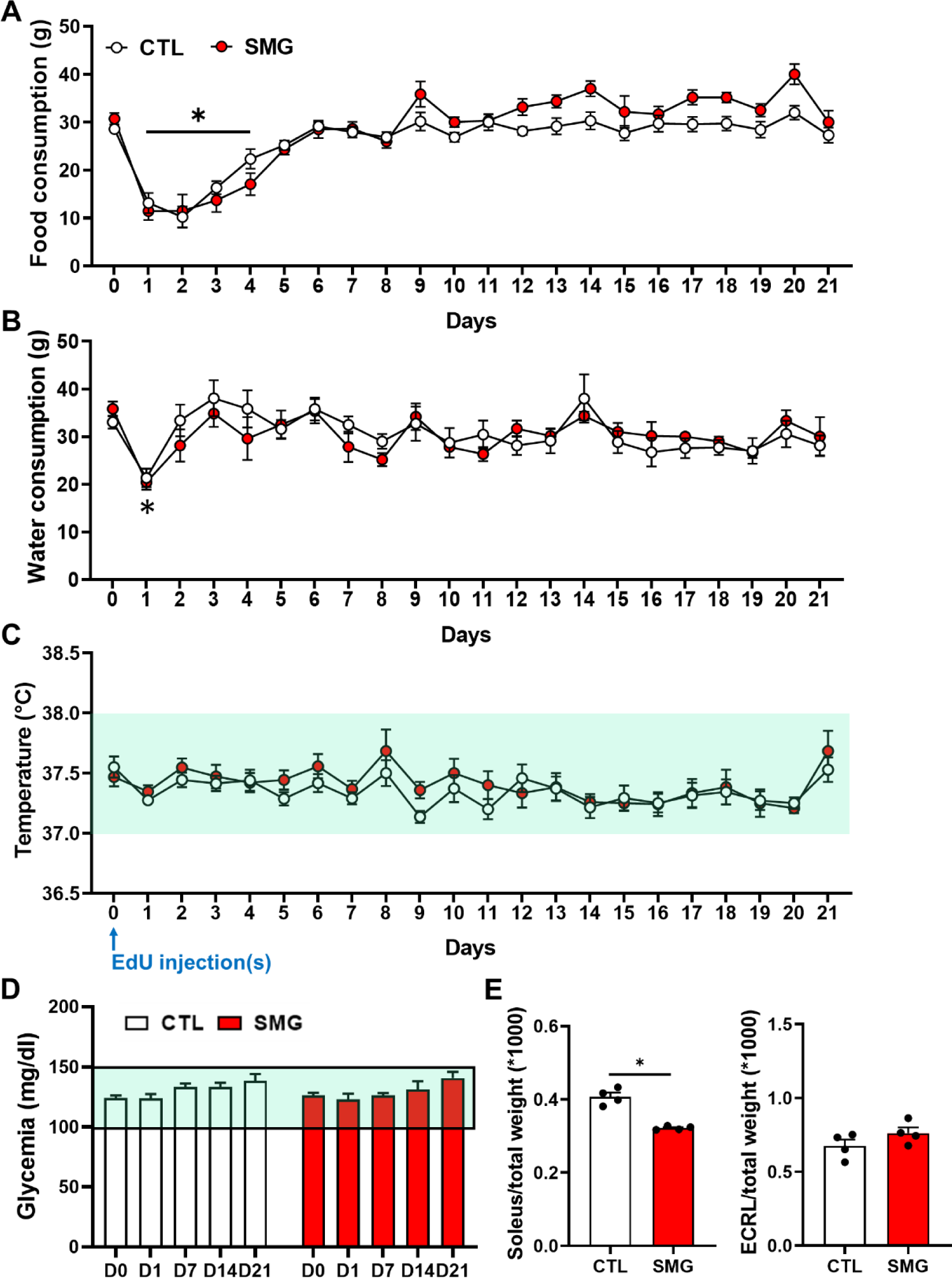
Physiological parameters of rats during the adult neurogenesis study. **A.** Food consumption of the CTL (white) and SMG (red) rats. **B.** Water consumption of the CTL (white) and SMG (red) rats. **C.** Temperature of the CTL (white) and SMG (red) rats. **D.** Glycemia of the CTL (white) and SMG (red) rats at D0, D1, D7, D14 and D21. Green area represents the physiological values. **E.** Weight of the soleus and ECRL muscles related to the total weight of the animals in CTL (white) and SMG (red) rats. All data are presented as mean ± SEM. * p < 0.05.

**Figure Sup. 2.**
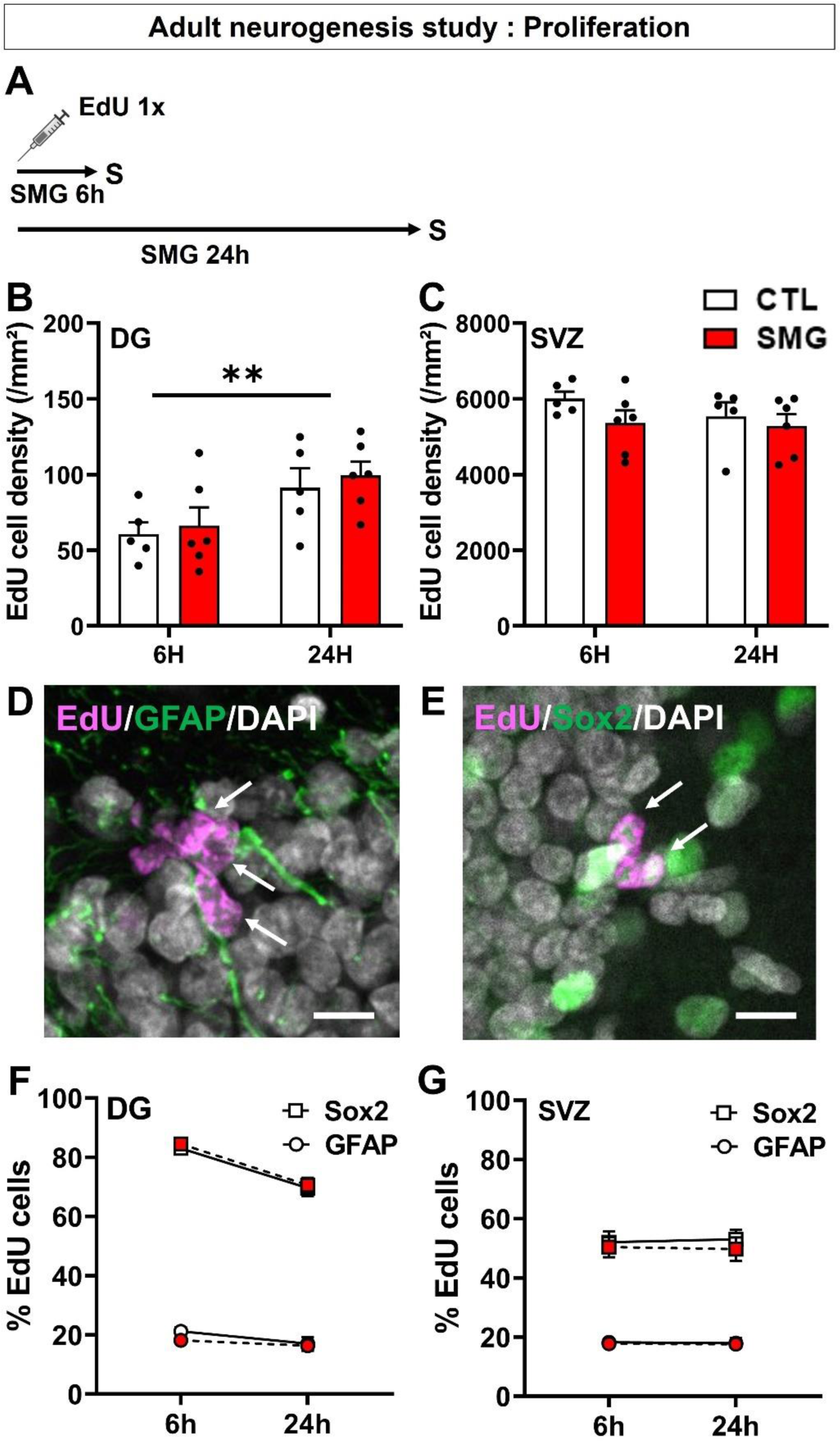
Newborn cells proliferation under SMG. **A.** Experimental design. Rats were injected with EdU and exposed to SMG during 6 or 24 hours. **B.** Density of EdU cells per mm² in the DG of CTL (white) and SMG (red) rats. **C.** Density of EdU cells per mm² in the SVZ of CTL (white) and SMG (red) rats. **D.** Example of EdU+ cells expressing GFAP in the DG of the hippocampus. Scale bar: 10µm. **E.** Example of EdU+ cells expressing Sox2 in the DG of the hippocampus. Scale bar: 10 µm. **F.** Percentage of EdU cells expressing GFAP or Sox2 in the DG of CTL (white) and SMG (red) rats. **G.** Percentage of EdU cells expressing GFAP or Sox2 in the SVZ of CTL (white) and SMG (red) rats. All data are presented as mean ± SEM. ** p < 0.01.

**Figure Sup. 3.**
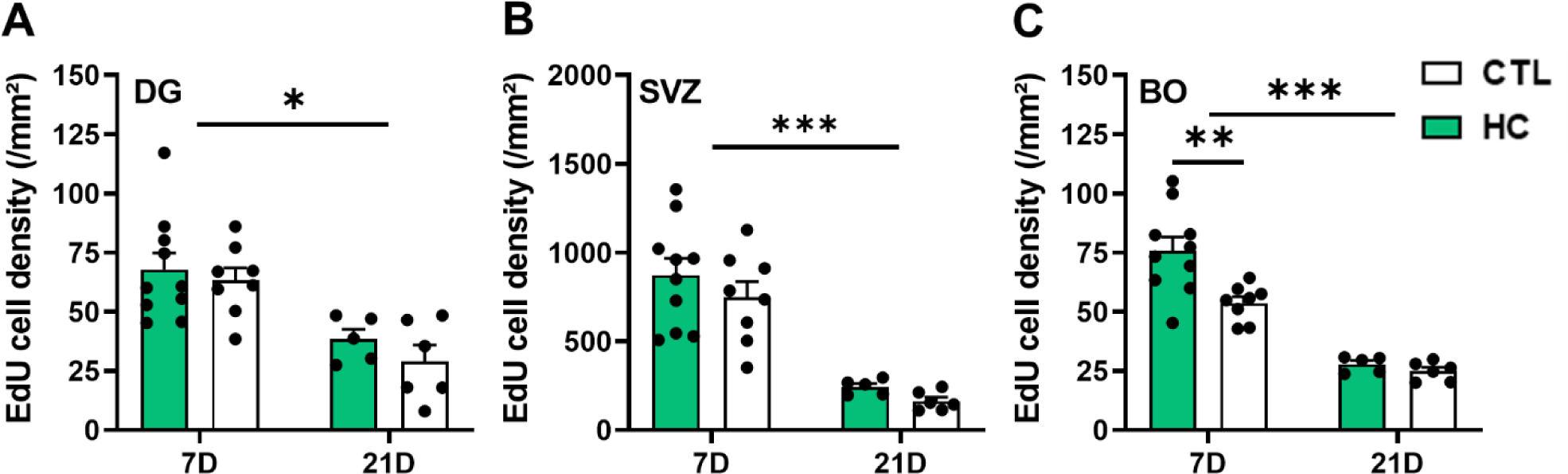
Newborn cells survival in single-housed (CTL) and grouped-housed rats (HC). **A.** Density of EdU cells per mm² in the DG of HC (green) and CTL (white) rats. No difference was observed between HC and CTL rats (two-way ANOVA, SMG effect F(1, 25) = 1.03, p = 0.32). **B.** Density of EdU cells per mm² in the SVZ of HC (green) and CTL (white) rats. No difference was observed between HC and CTL rats (two-way ANOVA, SMG effect F(1, 25) = 1.41, p = 0.25). **C.** Density of EdU cells per mm² in the OB of HC (green) and CTL (white) rats. A significant effect was observed between HC and CTL rats at D7 (two-way ANOVA, SMG effect F(1, 25) = 7.76, p = 0.01; Bonferroni multiple comparisons test: 7D p = 0.003, 21D p > 0.99). All data are presented as mean ± SEM. * p < 0.05, ** p < 0.01, *** p < 0.005.

**Figure Sup. 4.**
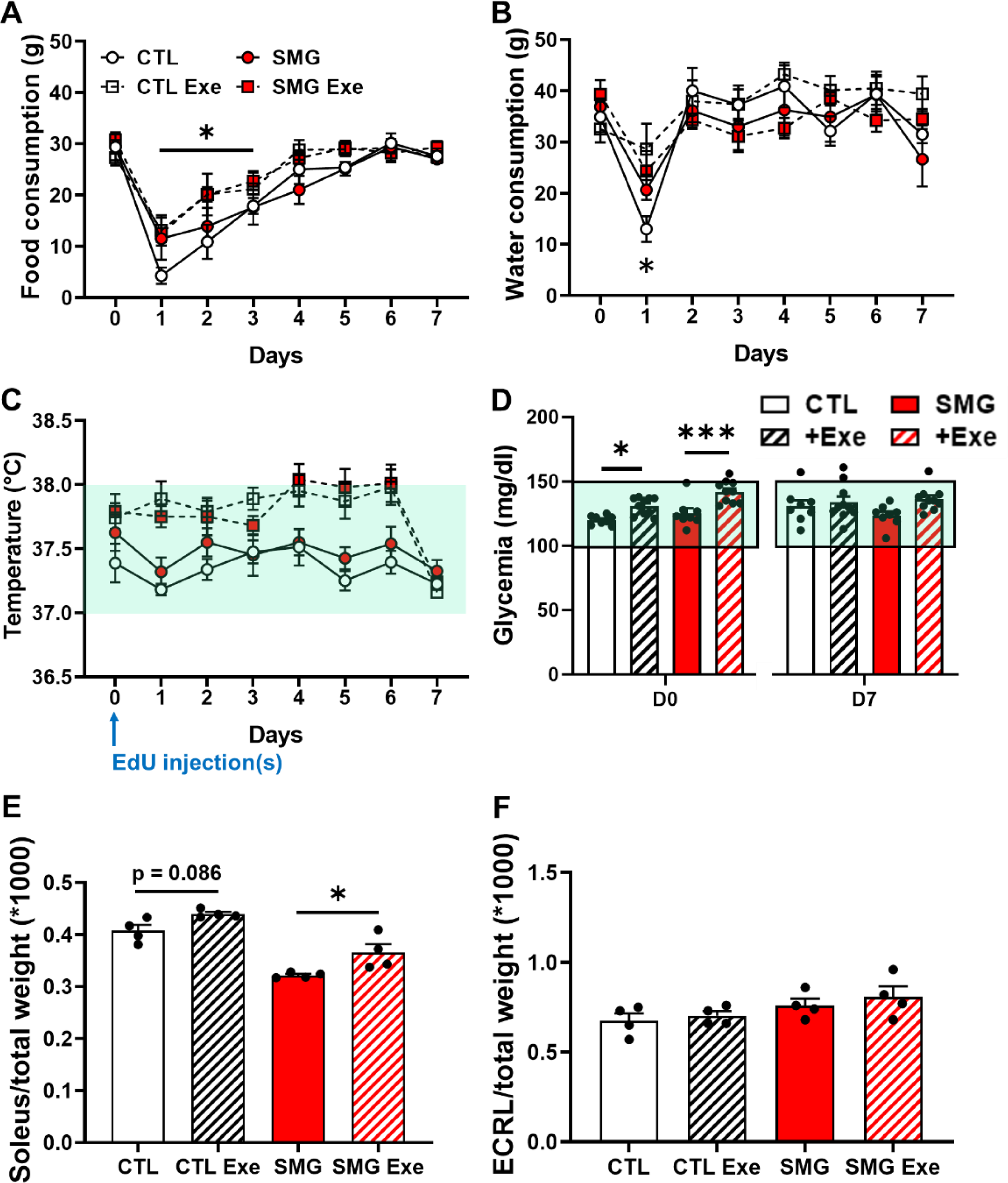
Physiological parameters of rats during the adult neurogenesis study with physical exercise. **A.** >Food consumption of the CTL (white) and SMG (red) rats exposed (scare) or not (circle) to physical exercise. **B.** Water consumption of the CTL (white) and SMG (red) rats exposed (scare) or not (circle) to physical exercise. **C.** Temperature of the CTL (white) and SMG (red) rats exposed (scare) or not (circle) to physical exercise. **D.** Glycemia of the CTL (white) and SMG (red) rats exposed or not to physical exercise at D0 and D7. **E.** Weight of the soleus muscle related to the total weight of the animals in CTL (white) and SMG (red) rats. **F.** Weight of the ECRL muscle related to the total weight of the animals in CTL (white) and SMG (red) rats. All data are presented as mean ± SEM. * p < 0.05.

